# Acute Pericentral Liver Injury Induces a Novel Transient Hepatocyte Population

**DOI:** 10.1101/2025.09.17.676911

**Authors:** Syeda Nayab Fatima Abidi, Kerstin Seidel, Ginny Xiaohe Li, Matthew Fernandez, Robert Piskol, Eliah R. Shamir, Christian W. Siebel, Louis Vermeulen

## Abstract

The liver exhibits robust regenerative capacity in response to injury, a property shaped by its essential metabolic and detoxification roles. While the mechanisms of regeneration following chronic liver injury have been extensively studied, the response during the early phases of acute injury remains poorly understood. Here, we employed a pericentral model of acute liver injury using carbon tetrachloride (CCl4) to investigate the initial hepatocyte response to damage. Through integration of single-nucleus multiome-sequencing, lineage tracing, spatial transcriptomics, and immunostaining, we identified a novel, transient population of damage-responsive SOX9+ pericentral hepatocytes. This population does not exhibit progenitor-like behavior but instead potentially mediates key aspects of early tissue repair. Specifically, SOX9+ pericentral hepatocytes engage in interactions with hepatic stellate cells and display migratory features associated with wound closure. Moreover, these cells exhibit an immune-related transcriptional signature and colocalize with liver macrophages at injury sites, suggesting a role in modulating immune responses. Our findings highlight a previously underappreciated function of SOX9 as a stress-responsive regulator of hepatocyte behavior and intercellular crosstalk during the early stages of acute liver injury.

## Introduction

The liver possesses a remarkable capacity for regeneration, a trait consistent with evolutionary adaptation, given its constant exposure to environmental insults and its essential role in metabolic regulation and detoxification. Histologically, the liver is divided into repeating functional units referred to as lobules. Each lobule forms a hexagon organized around a central vein in the middle, with six portal triads at the corners (composed of the portal vein, the hepatic artery and the bile duct) (Figure 1a). Hepatocytes are the main parenchymal cell of the liver, and while they appear morphologically similar across the lobule, they are functionally heterogenous and can be divided into arbitrary zones along the portal-central axis, based on gene expression and metabolic gradients (Jungermann and Keitzmann, 1996). The two primary functional compartments are the periportal (PP) and the pericentral (PC) zones (Jungermann and Katz, 1989). These can be further subdivided to include a “midlobular” zone between the PP and PC zones, or further refined into nine zonal layers (Colnot and Perret, 2010; Halpern et al., 2017).

**Figure 1.**
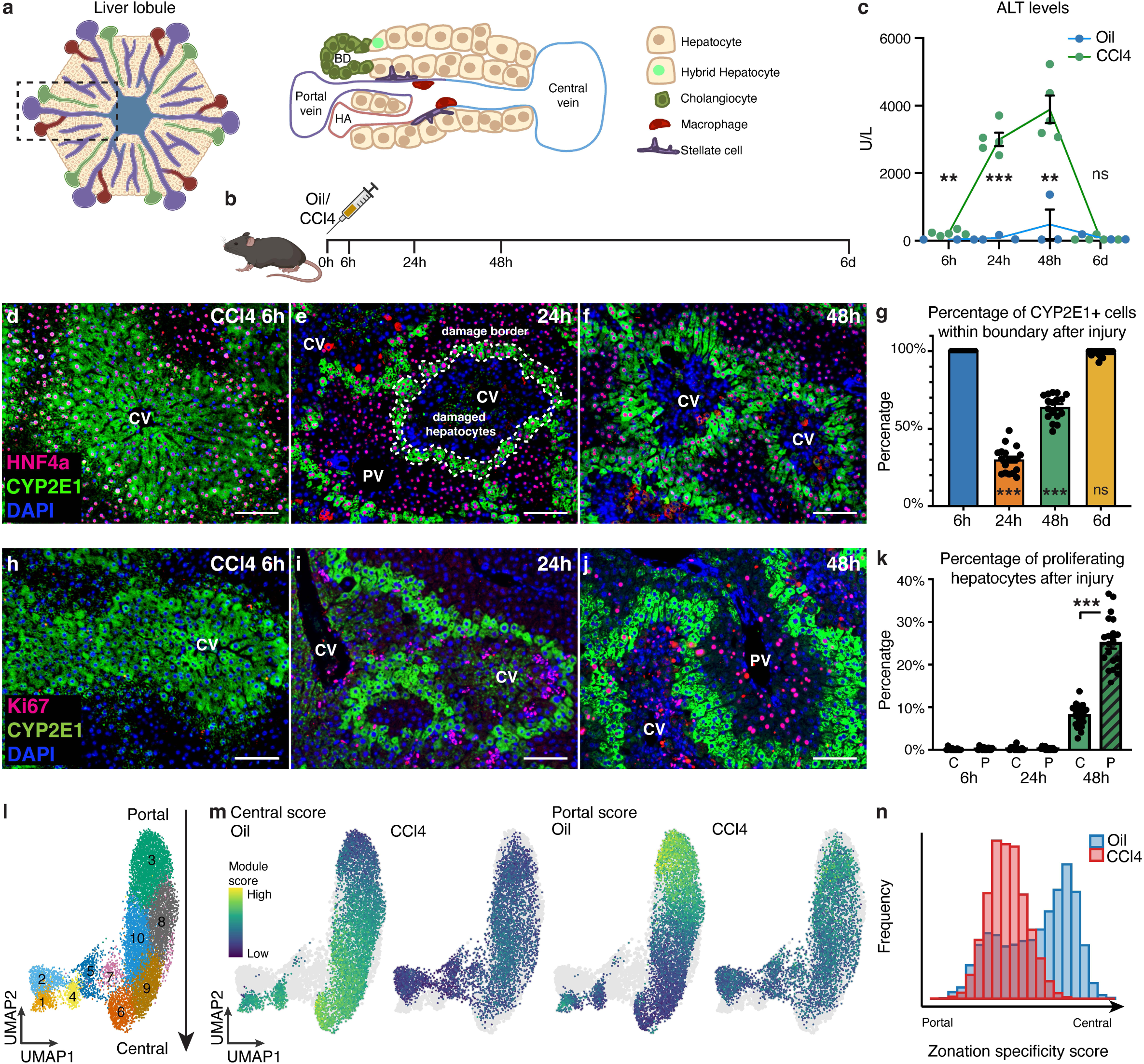
Early stages of damage and repair progression in CCl4-induced acute injury. (a) Schematic of a healthy liver lobule, showing different cell types present in the homeostatic state. Schematic adapted from (Gordillo et al., 2015). (b) Schematic of the experimental set-up. (c) Serum ALT levels at the indicated time points (n = 3 for oil, and n = 5 for CCl4, for each time point). p-values: 6h = 0.007, 24h = 3.51E-05, 48h =0.002, and 6d = 0.72. (d-f) Representative co-stain images for HNF4a and CYP2E1 in mice treated with CCl4 at 6h(d), 24h (e), and 48h (f) post treatment. CV = central vein, PV = portal vein. (g) Quantification of the number of CYP2E1+ nuclei as a percentage of the total number of nuclei found within the CYP2E1 boundary. p-values: 24h = 1.96E-21, 48h =4.23E-14, and 6d = 0.07. 24h, 48h and 6d compared against 6h for statistical analysis. (h-j) Representative co-stain images for Ki67 and CYP2E1 in mice treated with CCl4 at 6h(h), 24h (i), and 48h (j) post treatment. (k) Quantification of proliferating hepatocytes after injury in the PC and PP regions, at the indicated time points. The number of Ki67+/CYP2E1+ nuclei were quantified as a percentage of total CYP2E1+ nuclei to give the percentage of proliferating hepatocytes in the central (C) region. The number of Ki67+ nuclei outside the CYP2E1 boundary were quantified as a percentage of total nuclei outside the CYP2E1 boundary to give the percentage of proliferating hepatocytes in the portal (P) region. Comparison between portal and central proliferation – p-values: 6h = 0.30, 24h =0.96, and 48h = 1.47E-13. (l) UMAP of hepatocyte subset, from both oil and CCl4 samples. (m) Central and portal hepatocyte signature (Table 1) for the hepatocyte subset, split by oil and CCl4 samples. (n) Zonation specificity score for the oil and CCl4 samples. Zonation specificity scores were calculated as follows: the central and portal hepatocyte scores were scaled to a value between 0 and 1, these scaled values were subsequently used to calculate zonation specificity score using the formula – zonation specificity score = central score/(central score + portal score). Student’s T-test used for statistical analysis. Error bars represent SEM. Scale bars are 100 μm.

As detoxification of harmful substances is a key function of the liver, the metabolic zonation becomes important in the context of liver pathologies, with specific enzymes metabolizing their target toxins, resulting in zone-specific patterns of injury (Lindros, 1997). Given the clinical relevance of liver disease, it is important to have a detailed understanding of the liver regeneration process. Classically, liver regeneration has been studied using surgical models such as partial hepatectomy; however, increasingly, focus has shifted to studying pathologically relevant models, like toxin-mediated injury (Leventhal et al., 2020). Among these, chronic injury models have been extensively studied to explore mechanisms underlying disease conditions. In contrast, acute injury remains less well characterized. Acute liver failure (ALF) is a rare syndrome, with unexpected rapid onset of severe liver injury in otherwise healthy individuals, leading to a frequently fatal outcome. Common causes, like acetaminophen toxicity and viral hepatitis, can lead to ALF, and in severe cases, emergency liver transplantation is the only therapeutic option (Stravitz and Lee, 2019). Considering the high mortality rate, and the limited availability of donor organs and other therapeutic options, a greater focus on acute injury, especially in the early timepoints immediately following damage, will be important to improve outcomes for patients affected by this condition.

Numerous studies have established that the primary mechanism of hepatocyte recovery is by the proliferation of existing hepatocytes (Malato et al., 2011; Matsumoto et al., 2020; Schaub et al., 2014; Tarlow et al., 2014; Yanger et al., 2014). However, conflicting lineage-tracing studies have proposed that specific hepatocyte subsets may function as stem-like cells with enhanced regenerative potential (Ang et al., 2019; Bralet et al., 1994; Font-Burgada et al., 2015; Lin et al., 2018; Planas-Paz et al., 2016; Pu et al., 2016; Sun et al., 2020; Wang et al., 2015; ZAJICEK et al., 1985), leading to uncertainty about the true origin of proliferating hepatocytes. With improved tools being available to investigate this question, recent studies have employed multiple unbiased approaches to demonstrate that regenerative capacity is not confined to a specialized subset, but rather is broadly distributed across the hepatocyte population, and midlobular hepatocytes appear to have the highest proliferative advantage (Chen et al., 2020; He et al., 2021; Lin et al., 2023; Wei et al., 2021). While these insights represent important progress, the precise hepatocyte response during the earliest stages of acute liver injury remains incompletely understood, warranting further investigation.

In this study, we employed a PC model of acute injury using carbon tetrachloride (CCl4) to investigate the early hepatocyte response to damage. By initially characterizing the hepatocyte response at early timepoints, we discovered a novel, transient, *Sox9+* damage-responsive cell population. We further integrated single-nucleus multiome-sequencing, lineage-tracing, spatial transcriptomics, and immunostaining validation, to demonstrate that: (1) SOX9 is the predominantly active transcription factor in damaged hepatocytes; (2) the damage-responsive *Sox9+* cell population does not behave as a progenitor cell type; (3) this cell population interacts with stellate cells and potentially responds to migratory signals to mediate wound closure, similar to the previously-described ANXA2+ migratory hepatocytes (Matchett et al., 2024); and, most strikingly, (4) this cell population displays an immune signature and closely colocalizes with liver macrophages at the site of injury to potentially mediate their function. Thus, our data argue for SOX9 as a stress response mediator that helps mediate multiple facets of tissue repair, in addition to its traditionally accepted role as a progenitor marker.

## Results

### Characterization of early stages of damage and repair progression in CCl4-induced acute injury

To characterize the early hepatocyte response, we utilized the CCl4-induced model of acute liver injury. CCl4 metabolization by the CYP2E1 enzyme produces highly reactive radical species that damage cellular macromolecules to cause cell swelling, leakage of enzymes, and ultimately cell death (Knockaert et al., 2012; Scholten et al., 2015; Weber et al., 2003; Wong et al., 1998). As CYP2E1 is exclusively expressed in the PC half of the lobule (Halpern et al., 2017), hepatotoxicity is restricted to this compartment. Using this damage model, we treated 8-week-old mice with a single dose of CCl4 (damage induction) or corn oil (vehicle control), and then harvested the livers at 6h, 24h, 48h and 6 days after injury (Figure 1b). We observed morphological changes as early as 6h after damage induction (Figure S1a,b), with swollen hepatocytes appearing at 24h and persisting until 48h (Figure S1c,d). By 6 days after CCl4 administration, the lobule structure had returned to its normal morphological pattern (Figure S1e). We confirmed hepatocellular damage by measuring serum ALT (alanine aminotransferase) levels and found them to be elevated starting at 6h, perduring until 48h, and returning to baseline levels at 6 days (Figure 1c). Consistent with the histological data, CYP2E1 immunostaining showed that peak damage occurred at 24h after damage induction, leaving only 1-2 cell layers of CYP2E1+ hepatocytes at the PC border (Fig 1d-f, and Figure S1f,g). Using CYP2E1 expression as our guide, we divided the PC region into the CYP2E1+ damage border, and the area within this border as the damaged hepatocytes (Figure 1e). Intriguingly, while the CYP2E1+ hepatocytes were still positive for the hepatocyte marker HNF4a at 6h and 24h (Figure 1d,e), the damaged hepatocytes at 24h had almost completely lost HNF4a expression (Figure 1e). Histologically, these damaged hepatocytes were not fully necrotic at this timepoint (Figure S1c), which suggests a disruption in hepatocyte fate. However, the PC region was already beginning to recover by 48h, and was almost completely regenerated at 6 days, with restoration of CYP2E1 expression (Figure 1f,g, Figure S1g).

We next examined the proliferation response to damage. Within the PC region, we restricted our analysis to the CYP2E1+ hepatocytes, to avoid confounding our results due to the presence of other cell types and necrotic hepatocytes. We defined the CYP2E1-hepatocytes outside the CYP2E1+ border as PP hepatocytes. Consistent with other studies, the hepatocyte proliferation response was delayed to 48h (Figure 1h-k, and Figure S1h,i) (Matchett et al., 2024). Interestingly, the initial proliferation response was significantly higher in the PP region as opposed to the PC region at 48h (Figure 1k). This is consistent with findings from Wang and colleagues, which showed significantly higher PP hepatocyte proliferation 2 days after CCl4 injury, with PC hepatocyte proliferation increasing at later time points, with its peak at 3 days after injury (Wang et al., 2024). Consistent with histology and CYP2E1 expression data, proliferation was reduced to basal levels at 6 days (Figure S1j). Altogether, these data indicate that injury triggers early morphological changes and potential fate transitions, followed by a delayed proliferative response.

### Single-nuclear transcriptomics analysis to investigate early damage response

To further detail the early hepatocyte response, we performed single-nuclear profiling of gene expression (snRNA-seq) and chromatin accessibility (snATAC-seq) on undamaged (oil treated), and damaged (CCl4 treated) samples at 24h after treatment. This timepoint was selected to coincide with peak damage (Figure 1e), and to capture the earliest significant molecular responses. We began by exploring the snRNA-seq dataset using the ArchR package (15,245 nuclei) and obtained 12 total clusters for both samples (Figure S2a-c), which were annotated using known lineage markers (Figure S2d,e, and Table 1). While the mesenchymal, endothelial and immune cells from undamaged and damaged livers clustered together, the hepatocytes were strikingly divided between the two samples, indicating vastly distinct transcriptional profiles (Figure S2c). The large hepatocyte cluster showed a clear zonation pattern from central to portal for the undamaged sample (Figure S2f, and Table 1). To focus on the early hepatocyte response, we isolated the hepatocyte cluster for detailed exploration.

We recovered a total of 11,926 hepatocyte nuclei, that were subclustered into 10 new clusters (Figure 1l, and Figure S2g,h). While examining the hepatocyte lineage score, and central and portal zone scores, we noticed that the damaged hepatocytes displayed a considerably lower signature for these categories (Figure 1m, and Figure S2i). Plotting the zonation specificity score showed a bimodal distribution for the undamaged sample, representing both the PP and the PC hepatocytes, however, this distribution was lost in the damaged sample. The resulting distribution revealed an increase in hepatocytes exhibiting both portal and central characteristics, with a shift toward a more portal-like identity (Figure 1n). These data suggest a loss in the portal-central polarity of the lobule, in addition to the loss of PC cells, consistent with our damage model. Together with the observed loss of HNF4a in the damaged hepatocytes (Figure 1e), these findings suggest a broader disruption in hepatocyte identity and indicate potential functional plasticity among the remaining hepatocytes.

### Discovery of a novel SOX9+ cell population activated during early acute injury

Hepatocytes and cholangiocytes (biliary epithelial cells) originate from a common bi-potent hepatoblast population during liver development (Lemaigre and Zaret, 2004). A subset of hepatoblasts express the SOX9 transcription factor to specify cholangiocyte fate (Antoniou et al., 2009), and line up around the portal mesenchyme to form the ductal plate (Eyken et al., 1988). Intriguingly, the SOX9+ embryonic ductal plate cells retain the capacity to differentiate into both periportal hepatocytes as well as cholangiocytes (Carpentier et al., 2011). Additional studies have also described the existence of SOX9+ periportal hepatocytes (hybrid hepatocytes) in the homeostatic adult liver, that can give rise to both hepatocytes and cholangiocytes in response to chronic damage, but do not contribute significantly to acute injury repair (Font-Burgada et al., 2015; Han et al., 2019). It is important to note that these bipotential, progenitor-like cell populations in the liver exist in the specific periportal niche. Given SOX9’s association with progenitor identity, and its role in chronic liver repair, we sought to investigate its expression dynamics in our model of acute injury.

Consistent with previous studies, immunostaining for SOX9 showed expression restricted to cholangiocytes in the homeostatic undamaged liver (Figure 2a), and the rare hybrid hepatocytes present in the periportal niche (Figure S3a). Remarkably, we discovered a novel SOX9+ hepatocyte population activated in the damaged PC region (Figure 2b-d). At 6h after damage induction, the SOX9+ hepatocytes co-expressed the hepatocyte marker HNF4a (Figure 2b,c), similar to the hybrid hepatocytes, however, they had lost the HNF4a expression by 24h (Figure 2d). To quantify this damage-responsive SOX9+ cell population, we co-stained SOX9 with CYP2E1 to define the PC region (Figure 2e-h). While SOX9+/CYP2E1+ cells were never present in the undamaged liver (Figure 2e), they appeared in large numbers at 6h after damage induction and had increased significantly by 24h (Figure 2h). Interestingly, the appearance of these SOX9+ cells was transient in the damaged PC region as very few SOX9+/CYP2E1+ were found at 48h (Figure 2h, and Figure S3b), and SOX9 expression had returned to a normal pattern by day 6 (Figure S3c). Conversely, a second SOX9+ hepatocyte population was observed at 48h, now in the PP region (Figure S3b). As the PP niche already consists of some homeostatic SOX9+ hepatocytes, we quantified PP SOX9+ nuclei, and found that these were significantly increased after damage at 48h (Figure S3d). Since we had observed an enriched PP proliferative response at 48h (Figure 1j,k), we examined the proportion of proliferating SOX9+ nuclei at 48h (Figure S3e-h). While a percentage of PP SOX9+ nuclei proliferate (Figure S3f-h), the majority of proliferating nuclei at this time point were not SOX9+ (Figure S3f,g). We speculate that this second SOX9+ PP hepatocyte population is the same as the hybrid hepatocytes, which were reported to only make a minor contribution to acute injury (Font-Burgada et al., 2015). Notably, the early damage-responsive SOX9+ population observed at 6h and 24h, differs from previously characterized SOX9+ hepatocyte populations, as it is found in the PC region rather than the traditional PP niche.

**Figure 2.**
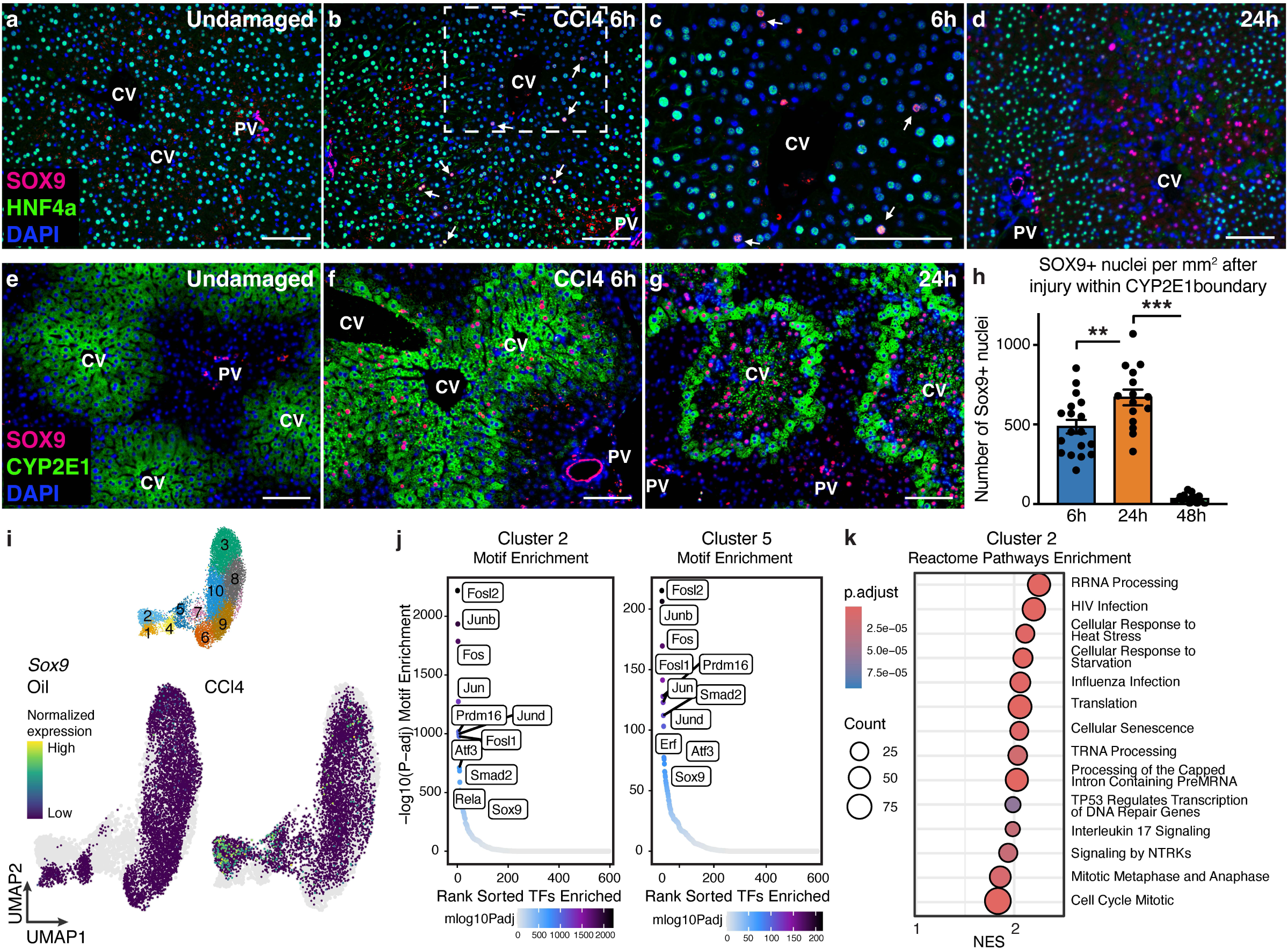
Discovery of a novel SOX9+ cell population activated during early acute injury. (a-d) Representative co-stain images for SOX9 and HNF4a in mice treated with oil (a) or CCl4, 6h (b,c), and 24h (d) post treatment (n = 3 for oil, and n = 5 for CCl4, for each time point). Arrows represent SOX9+/HNF4a+ hepatocytes. (e-g) Representative co-stain images for SOX9 and CYP2E1 in mice treated with oil (e) or CCl4, 6h (f), and 24h (g) post treatment. (h) Quantification of the number of SOX9+ nuclei within the area of the CYP2E1+ boundary. p-values: 6h-24h = 0.008, 24h-48h = 2.65E-14. (i) UMAP showing *Sox9* expression for the hepatocyte subset, split by oil and CCl4 samples. (j) Motif enrichment in differential peaks for hepatocyte subset cluster 2, and cluster 5. (k) Reactome Pathways Enrichment for the hepatocyte subset cluster 2. Student’s T-test used for statistical analysis. Error bars represent SEM. Scale bars are 100 μm.

We next leveraged our single-nucleus multiome data to further investigate the identity and role of these newly identified damage-responsive SOX9+ PC hepatocytes. Consistent with our immunostaining data, *Sox9* expression was confined to the damaged hepatocyte clusters and the PP hepatocytes (Figure 2i, and Figure S3i). Clusters 2 and 5 from the hepatocyte subset were significantly enriched for *Sox9* expression (Figure 2i). Consistent with its transcriptional upregulation, SOX9 also emerged as one of the most significantly enriched motifs in these clusters based on our snATAC-seq data (Figure 2j, Table 2). The motif enrichment also identified the NF-κB family member *Rela* (NF-κB p65), which has previously been shown to directly regulate *Sox9* induction by binding to its promoter (Figure 2j, Table 2) (Ushita et al., 2009). The other enriched motifs represented stress response genes like *Fos, Fosl1, Fosl2, Jun, Junb, Jund* and *Atf3*. Gene Set Enrichment Analysis (GSEA) for cluster 2 also revealed pathways related to stress response, but interestingly also showed enrichment of pathways related to cell cycle and mitosis (Figure 2k, and Table 3). These findings suggest that a subset of the damage-responsive SOX9+ PC hepatocytes is predominantly engaged in the immediate stress response following injury, activating pathways involved in DNA damage repair, but also indicate that these cells may have a proliferative advantage.

### Lineage tracing the damage-responsive SOX9+ cells reveals non-progenitor behavior

The association of SOX9 with bipotential hepatobiliary cells, along with our data suggesting a possible proliferative advantage, motivated us to investigate whether the early damage-responsive SOX9+ PC hepatocytes could contribute to acute injury repair, analogous to the role of hybrid hepatocytes in chronic liver injury. To this end, we used the transgenic *Sox9-CreER^T2^* mouse line (Kopp et al., 2011) (Figure S4a) crossed to a *Rosa26-LSL-tdTomato* reporter. Unexpectedly, despite multiple attempts and adjustments to the timing of tamoxifen administration, this line failed to label the damage-responsive SOX9⁺ PC hepatocytes (Figure S4b-d). To confirm that this line still activated *Sox9* expression in response to damage, we examined *Sox9* mRNA. Indeed, *Sox9* mRNA was observed in the PC region as early as 6h after damage (Figure S4e). To explain the failure of labeling, we also examined *Cre* mRNA expression. Interestingly, while *Cre* mRNA was observed in the PP region (in the cholangiocytes and the hybrid hepatocytes), *Cre* was not expressed in the PC region (Figure S4f). We hypothesized that the transgenic *Sox9-CreER^T2^* line lacks critical regulatory elements necessary for activating damage-responsive *Sox9* expression. To address this limitation, we generated a knock-in inducible *Sox9-CreER^T2^*allele, designed to retain all endogenous regulatory elements, and enable more faithful *Cre* activation across all *Sox9*-expressing cells (Figure S4g). Consistent with the known SOX9 expression pattern, our knock-in line crossed to the *Rosa26-LSL-tdTomato* reporter (Figure 3a), only labeled cholangiocytes and PP hepatocytes under homeostatic conditions (Figure S4h). Remarkably, while tamoxifen administration 24h prior to oil or CCl4 treatment labeled the cholangiocytes and PP hepatocytes, it was still unable to label the majority of the damage-responsive SOX9+ PC hepatocytes (Figure S4i,j). These results demonstrate that *Sox9* is induced *de novo* in response to injury, and confirm our original hypothesis that the damage-responsive SOX9+ PC hepatocytes represent a distinct cell population, rather than an expansion of a pre-existing SOX9+ hepatocyte subset.

**Figure 3.**
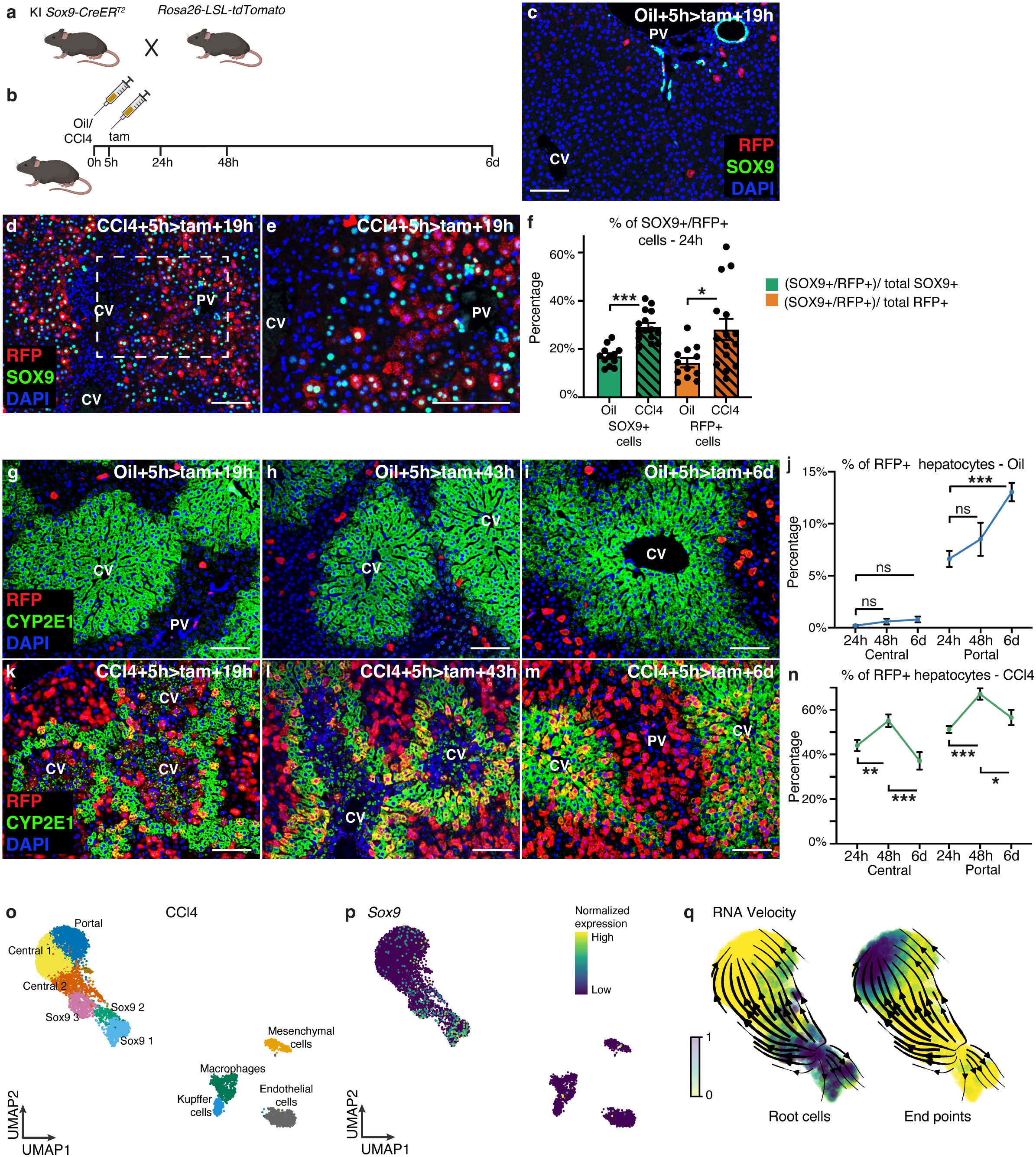
Lineage tracing the damage-responsive *Sox9+* cells reveals non-progenitor behavior. (a) Schematic of the mouse lines used for performing the lineage tracing experiments. (b) Schematic of the experimental set-up. (c) Representative co-stain image for RFP and SOX9 in mice treated with oil (n = 3) at 24h post treatment. (d,e) Representative co-stain images for RFP and SOX9 in mice treated CCl4 (n = 4) at 24h post treatment. (f) Quantification of SOX9+/RFP+ dual positive cells as a percentage of total SOX9+ cells, and total RFP+ cells at 24h, for oil and CCl4 treatment. (g-i) Representative co-stain images of RFP and CYP2E1 at 24h (g), 48h (h), and 6d (i) post oil treatment. (n = 3 for each time point). (j) Quantification of the percentage of RFP+ hepatocytes in the central and the portal regions at the indicated time points after oil treatment. Comparison for central region – p-values: 24h-48h = 0.17, 24h-6d = 0.03, comparison for portal region – p-values: 24h-48h = 0.29, 24h-6d = 8.85E-06. (k-m) Representative co-stain images of RFP and CYP2E1 at 24h (k), 48h (l), and 6d (m) post CCl4 treatment. (n = 4 for each time point). (n) Quantification of the percentage of RFP+ hepatocytes in the central and the portal regions at the indicated time points, after CCl4 treatment. Comparison for central region – p-values: 24h-48h = 0.006, 48h-6d = 0.0006, comparison for portal region – p-values: 24h-48h = 3.99E-06, 48h-6d = 0.02. (o) UMAP of cell lineages based on known lineage signatures, from just the damaged sample. (p) UMAP showing *Sox9* expression for the damaged sample only. (q) RNA velocity analysis of the damaged sample, showing the root cells and end points (terminal state). Student’s T-test used for statistical analysis. Error bars represent SEM. Scale bars are 100 μm.

We reasoned that tamoxifen administration after damage induction would give us the highest chances of capturing the *de novo Sox9* expression in PC hepatocytes (Figure 3b). Tamoxifen administration 5h after oil treatment only labeled the cholangiocytes and PP hepatocytes, as observed previously (Figure 3c). Strikingly, CCl4 treatment led to a marked increase in RFP labeling across the entire lobule at 24h (Figure 3d). We also observed SOX9 expression across the lobule, including the PP region, explaining the RFP labeling (Figure 3d,e). We quantified the number of dual SOX9+/RFP+ cells as a percentage of total SOX9+ cells, and of total RFP+ cells at 24h (Figure 3f). For oil treatment, the dual positive cells form ∼17% of all SOX9+ cells (including cholangiocytes and PP hepatocytes). Surprisingly, the dual positive cells only form ∼14% of all RFP+ cells, with the majority of RFP+ labeled hepatocytes being negative for SOX9 expression (Figure 3c,f). This result indicates that SOX9 expression may be transient in the PP hepatocytes. For CCl4 treatment, the proportion of the dual positive cells was significantly higher for both total SOX9+ and total RFP+ cells (Figure 3f). However, the majority of the RFP+ cells were still SOX9-negative, supporting the hypothesis of transient SOX9 expression. Accordingly, SOX9 expression had returned to baseline levels by day 6, consistent with morphological recovery seen at this timepoint (Figure S4k,l).

Since RFP labeling extended to the PP region, we co-stained with CYP2E1 to mark the PC region. In a time-course for the undamaged tissue, RFP labeled cells were largely absent in the PC region, only showing a minimal presence in the later time points (Figure 3g-j). The proportion of RFP+ labeled PP hepatocytes showed a gradual increase over time, likely reflecting a combination of sustained tamoxifen-induced labeling, and proliferation (Figure 3g-j). For the damaged PC region, we restricted our analysis to CYP2E1+ cells, to minimize confounding effects from other cell types and necrotic hepatocytes. Intriguingly, the proportion of RFP+ cells in both the PC and PP regions in the damaged tissue, increased from 24h to 48h, but declined significantly by day 6 (Figure 3k-n, and Figure S4m). This final decline in proportion argues against a progenitor-like function for these damage-responsive SOX9+ cells (Figure 3n).

We next performed RNA velocity analysis to determine whether transcriptional dynamics would support our lineage tracing findings. To specifically assess transcriptional trajectories within the damaged tissue, we subset our snRNA-seq data to isolate all the cells in the damaged sample (Figure 3o). For the damaged sample alone, there were 3 clusters that were enriched for *Sox9* expression which we labeled Sox9 1-3 (Figure 3o,p). RNA velocity analysis revealed a trajectory that originated from the Sox9 clusters and converged on PC hepatocytes as terminal states (Figure 3q and S4n). This is consistent with our lineage tracing data, which demonstrated that the damage-responsive SOX9+ PC hepatocytes do contribute to a proportion of the regenerated PC region (Figure 3m). Intriguingly, in the RNA velocity analysis, a subset of the Sox9-cells diverged from the main trajectory, suggesting an alternate fate (Figure 3q). Together, the lineage tracing and RNA velocity results suggest that the SOX9+ PC hepatocytes may support tissue repair through alternative mechanisms, rather than functioning as classical progenitor cells.

### *Sox9+* cells potentially mediate repair by interacting with stellate cells

Having ruled out a progenitor-like function for the SOX9+ PC hepatocytes, we turned back to our single-nucleus multiome data to explore their potential function in the injury response. We leveraged our paired RNA and ATAC data to infer transcription factor activity, and identified SOX9 as one of the most differentially active transcription factors in the two *Sox9*-enriched clusters (clusters 2 and 5) (Figure 2i, and Figure 4a), highlighting its potential functional importance. Consistently, clusters 2 and 5 were also enriched for *Sox9* target genes (Figure 4b). Interestingly, we observed that one of the most enriched target genes of *Sox9* was *Anxa2* (Figure 4b, and Figure S5a). Recently, an ANXA2+ migratory hepatocyte subpopulation has been described that emerges during liver regeneration, and plays a crucial role in effective wound closure (Matchett et al., 2024). We overlaid the “migratory hepatocyte” gene signature identified by Matchett and colleagues (Table 4) on our dataset, and found it to also be enriched in clusters 2 and 5 (Figure S5b). Using CYP2E1 protein expression to examine hepatocyte shape at 48h, we found that the hepatocytes closest to the necrotic region displayed motile morphology with extended lamellipodia, as opposed to the hepatocytes at the outer edge of CYP2E1 expression (Figure 4c,d). We also re-examined our lineage tracing data and found that a large proportion of these “motile” hepatocytes were RFP+, indicating that they were descendants of the damage-responsive SOX9+ hepatocytes (Figure 4e). These data suggest that one of the functions of the damage-responsive SOX9+ PC hepatocytes could be facilitating wound closure by regulating *Anxa2*.

**Figure 4.**
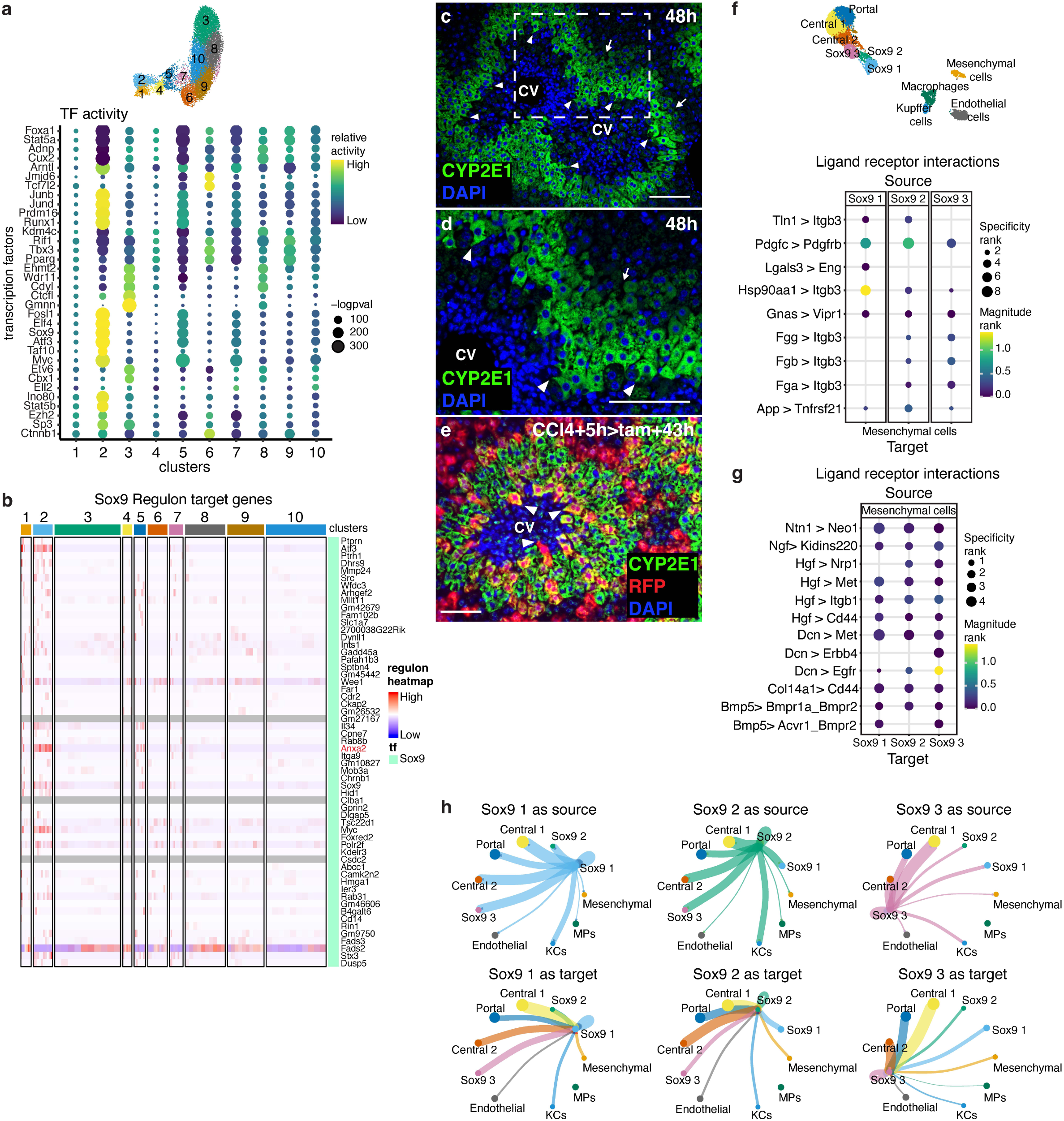
*Sox9+* cells potentially mediate repair by interacting with stellate cells. (a) Transcription Factor (TF) activity inferred using epiregulon, for all the clusters in the hepatocyte subset. (b) Sox9 Regulon target gene expression across all the clusters in the hepatocyte subset. (c,d) Representative IF image for CYP2E1, 48h after CCl4 treatment. Arrowheads indicate motile hepatocytes, while the arrows show normal hepatocyte morphology. (e) Representative co-stain images for CYP2E1 and RFP, with tam administration 5h post CCl4 injection. Arrowheads indicate motile hepatocytes. (f) Dot plot showing the ligand-receptor interactions between the Sox9 clusters (source) and mesenchymal cells (target). (g) Dot plot showing the ligand-receptor interactions between the mesenchymal cells (source) and Sox9 clusters (target). (h) Circle plot showing the total interaction strength between the Sox9 clusters and other cell-type clusters. Scale bars are 100 μm.

Interactome analysis has shown that ANXA2+ migratory hepatocytes potentially interact with mesenchymal cells (Matchett et al., 2024). To perform ligand-receptor interaction analysis on our data, we focused on the damaged sample to avoid any artificial interactions between the damaged and the undamaged sample. This analysis revealed multiple interactions between the Sox9 clusters and the mesenchymal cells (Figure 4f,g). Defining the Sox9 clusters as source cells and mesenchymal cells as targets, resulted in interactions important for Hepatic Stellate Cell (HSC) activation (Figure 4f) (Jiang et al., 2012; Kocabayoglu et al., 2015). Importantly, defining the mesenchymal cells as source cells, highlighted multiple interactions related to Hepatocyte Growth Factor (HGF), especially through its receptor Met, which has been shown to be critical for hepatocyte migration and wound closure (Figure 4g) (Huh et al., 2004; Matchett et al., 2024). Plotting overall interactions between the Sox9 clusters and all other cell types revealed that interactions were stronger when mesenchymal cells targeted the Sox9 clusters, compared to when Sox9 clusters targeted the mesenchymal cells (Figure 4h).

Since our data reflected important interactions between the Sox9 cluster cells and HSCs, we next examined the dynamics of HSCs at the early time points after damage induction (Figure S5c-g). In the undamaged liver, the HSCs were dispersed throughout the liver lobule (Figure S5c). This distribution did not change appreciably until 24h after CCl4-induced damage, when the HSCs began to organize close to the central veins (Figure S5d,e). By 48h, they had completely invaded the injury site, and now also expressed α-SMA, a marker for activated HSCs (Figure S5f). Consistent with morphological recovery, the HSC distribution had largely returned to normal by day 6, although a few HSCs remained localized near the central vein (Figure S5g). The mobilization of HSCs at 24h aligned with the peak SOX9 expression in the PC region. To determine whether the damage-responsive SOX9+ PC hepatocytes interact with HSCs, we co-stained for SOX9 and the HSC marker Desmin (Figure S5h-j). Notably, we found that HSCs were preferentially localized near SOX9 expressing cells, with a marked absence in regions lacking SOX9 expression (Figure S5i,j). Collectively, these data suggest that the damage-responsive SOX9+ PC hepatocytes not only have the potential to activate HSCs but, more crucially, they may respond to growth signals such as HGF from HSCs. This response could enable them to adopt a migratory phenotype, which is important in facilitating wound closure.

### *Sox9+* cells display immune signatures and interact with macrophages at the site of injury

The interactome plots for the Sox9 clusters (Figure 4h) revealed that their strongest interactions were with other hepatocytes, particularly the remaining PC hepatocytes. This would be consistent with the idea that these damage-responsive SOX9+ PC hepatocytes act as intermediaries – receiving signals from mesenchymal cells and relaying them to neighboring hepatocytes to coordinate effective wound closure. However, we also noticed strong interactions between the Sox9 clusters and myeloid cells (Figure 4h). To further explore this interaction, we performed GSEA on the remaining Sox9-expressing cluster (cluster 5) in the hepatocyte subset. This analysis revealed that the pathways were predominantly associated with TGF-β signaling, immune response signaling, and apoptosis (Figure 5a, and Table 5). TGF-β signaling has previously been shown to influence macrophage activity (Ashcroft, 1999; Patel et al., 2022; Zhang et al., 2016). This, along with the enrichment of pathways such as interferon signaling, interleukin signaling, and Toll-like receptor cascades, supports our initial hypothesis regarding interactions between the Sox9 clusters and myeloid cells.

**Figure 5.**
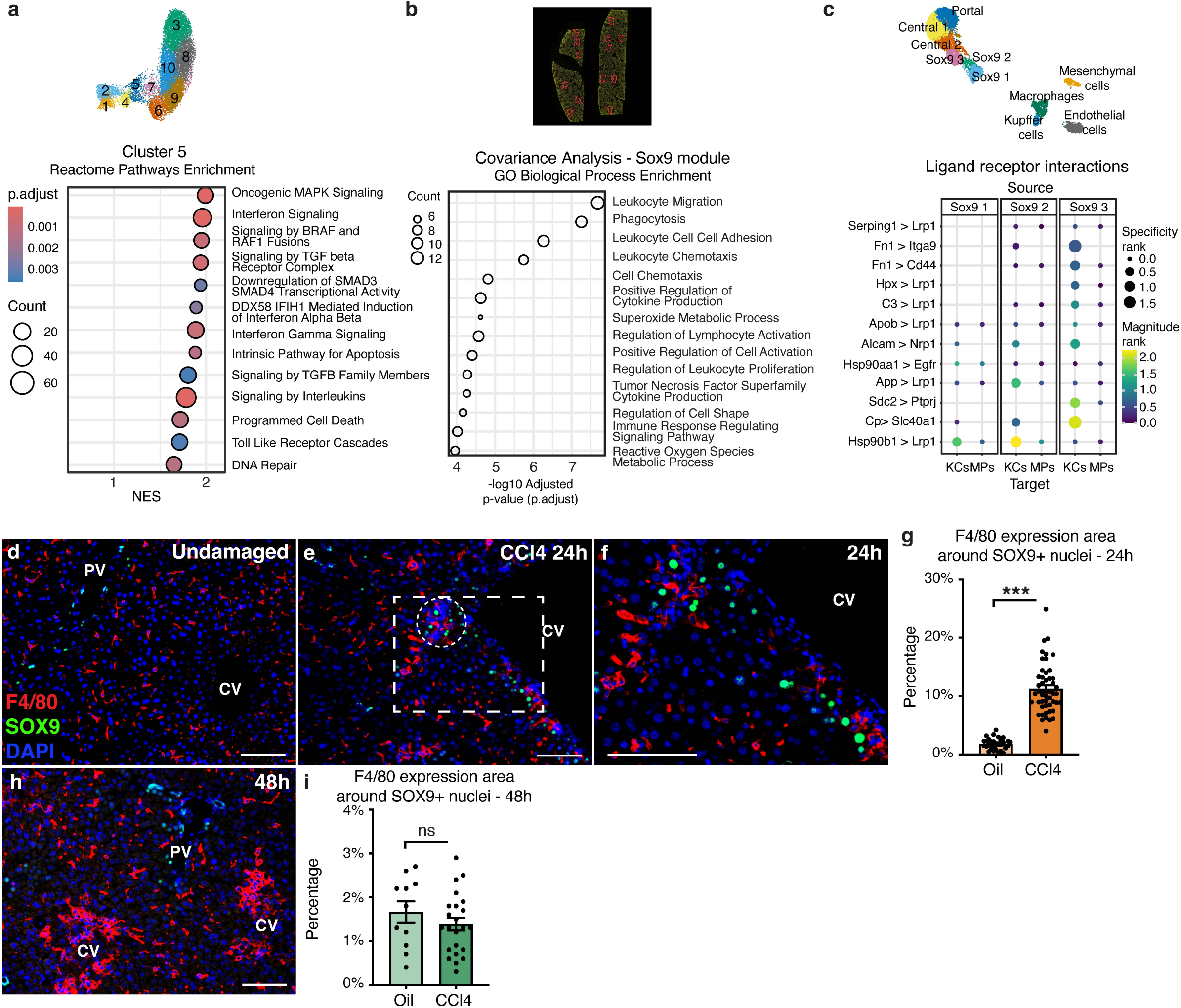
*Sox9+* cells display immune signatures and interact with macrophages at the site of injury. (a) Reactome Pathways Enrichment for the hepatocyte subset cluster 5. (b) GO Biological Process Enrichment for the Sox9 module identified via *WGCNA* from the GeoMx data. (c) Dot plot showing the ligand-receptor interactions between the Sox9 clusters (source) and myeloid cells (target), for the damaged sample only. (d-f) Representative co-stain images for F4/80 and SOX9 in mice treated with oil (d) or CCl4 at 24h (e,f) post treatment. (g) Quantification of the percentage of area covered by F4/80 expression in the vicinity of SOX9+ hepatocytes at 24h post treatment. A circle of a 100um diameter (∼5 hepatocytes) was drawn around the SOX9-expressing hepatocytes for calculating the percentage. p-value: 2.39E-19. (h) Representative co-stain image for F4/80 and SOX9 in mice treated with CCl4 at 48h post treatment. (i) Quantification of the percentage of area covered by F4/80 expression in the vicinity of SOX9+ hepatocytes at 48h post treatment. p-value: 48h = 0.31. Student’s T-test used for statistical analysis. Error bars represent SEM. Scale bars are 100 μm.

To further investigate the potential interaction between the damage-responsive SOX9+ PC hepatocytes and myeloid cells, we performed spatially restricted transcriptomic analysis using the GeoMx platform, at 24h after oil or CCl4 treatment. Using CYP2E1 expression as a marker, we defined multiple regions of interest (ROIs) per tissue section (Figure S6a), enabling selective RNA isolation from either undamaged PC hepatocytes, or damaged hepatocytes located within the CYP2E1+ damage border (Figure S6b). Principal Component Analysis (PCA) revealed clear distinctions between the undamaged and damaged ROIs (Figure S6c). Gene expression profiles further validated the earlier immunostaining and single-nucleus sequencing findings, demonstrating reduced *Cyp2e1* and elevated *Sox9* expression in the damaged ROIs (Figure S6d). GSEA also confirmed our earlier enrichment findings from the hepatocyte subset in the snRNA-seq data (Figure 2k, and Figure 5a), highlighting pathways involved in stress response, cell death, mitosis, and immune signaling (Figure S6e, and Table 6).

To specifically understand the transcriptional profile associated with the damage-responsive *Sox9*-expressing cells, we performed covariance analysis on the damaged ROIs. This analysis identified a *Sox9*-associated gene module, which was tested for over-representation against the GO Biological Process pathways. Strikingly, the results highlighted hallmark processes of immune activation, inflammation, and host defense (Figure 5b, and Table 7). Ligand–receptor interaction analysis from the snRNA-seq damaged sample, further identified multiple interactions between the Sox9 clusters and myeloid cells, highlighting pathways associated with cell adhesion, migration, and clearance of cellular debris (Figure 5c)(Boucher and Herz, 2011; Elpek, 2015a, 2015b; Gonias and Campana, 2014; Guo et al., 2019; Hight-Warburton and Parsons, 2019; Imhof and Aurrand-Lions, 2004; Lillis et al., 2008; Whiteford et al., 2011). Altogether, these data suggest that the damage-responsive SOX9+ PC hepatocytes may facilitate myeloid cell adhesion and retention at sites of injury, in addition to contributing to their activation upon arrival.

To further validate these findings, we examined the dynamics of liver macrophages at the early time points after damage induction (Figure S6f-j). Liver macrophages were distributed throughout the undamaged liver lobule (Figure S6f). Similar to HSC behavior, the macrophage distribution did not appreciably change until 24h, when clusters of macrophages could be seen around the central vein (Figure S6g,h), and by 48h, the macrophages had fully infiltrated the injury site (Figure S6i). Consistent with morphological recovery, macrophage distribution had returned to normal by day 6 (Figure S6j). The clustering of macrophages at 24h coincided with the peak damage-responsive SOX9 expression in the PC region, further indicating potential crosstalk between these populations in the early injury response. To determine whether the damage-responsive SOX9+ PC hepatocytes interact with macrophages, we co-stained for SOX9 and the macrophage marker F4/80 (Figure 5d-i). We observed a striking spatial association between macrophage clusters and the SOX9+ PC hepatocytes at 24h (Figure 5e-g), whereas no significant colocalization was observed with the SOX9+ PP hepatocytes at 48h (Figure 5h,i). Given that SOX9 is one of the most differentially active transcription factors in the SOX9+ PC hepatocytes, our findings reveal a potential non-progenitor function for SOX9 as a key mediator of the hepatocyte stress response, and as a regulator of myeloid cell engagement during the early phase of acute liver injury.

## Discussion

Based on our findings, we propose that a novel population of damage-responsive SOX9+ pericentral hepatocytes coordinates an early damage response in acute liver injury by interacting with HSCs and myeloid cells (Figure 6). Rather than serving a progenitor-like role, the SOX9+ population appears to function primarily as a mediator of stress response in the early phases of liver injury. In this capacity, SOX9 regulates key reparative processes, including hepatocyte migration and wound closure. Notably, our findings reveal an unexpected role for SOX9 in modulating myeloid cell recruitment and activity, suggesting that it is an important orchestrator of immune–parenchymal crosstalk during the initial stages of tissue repair.

**Figure 6.**
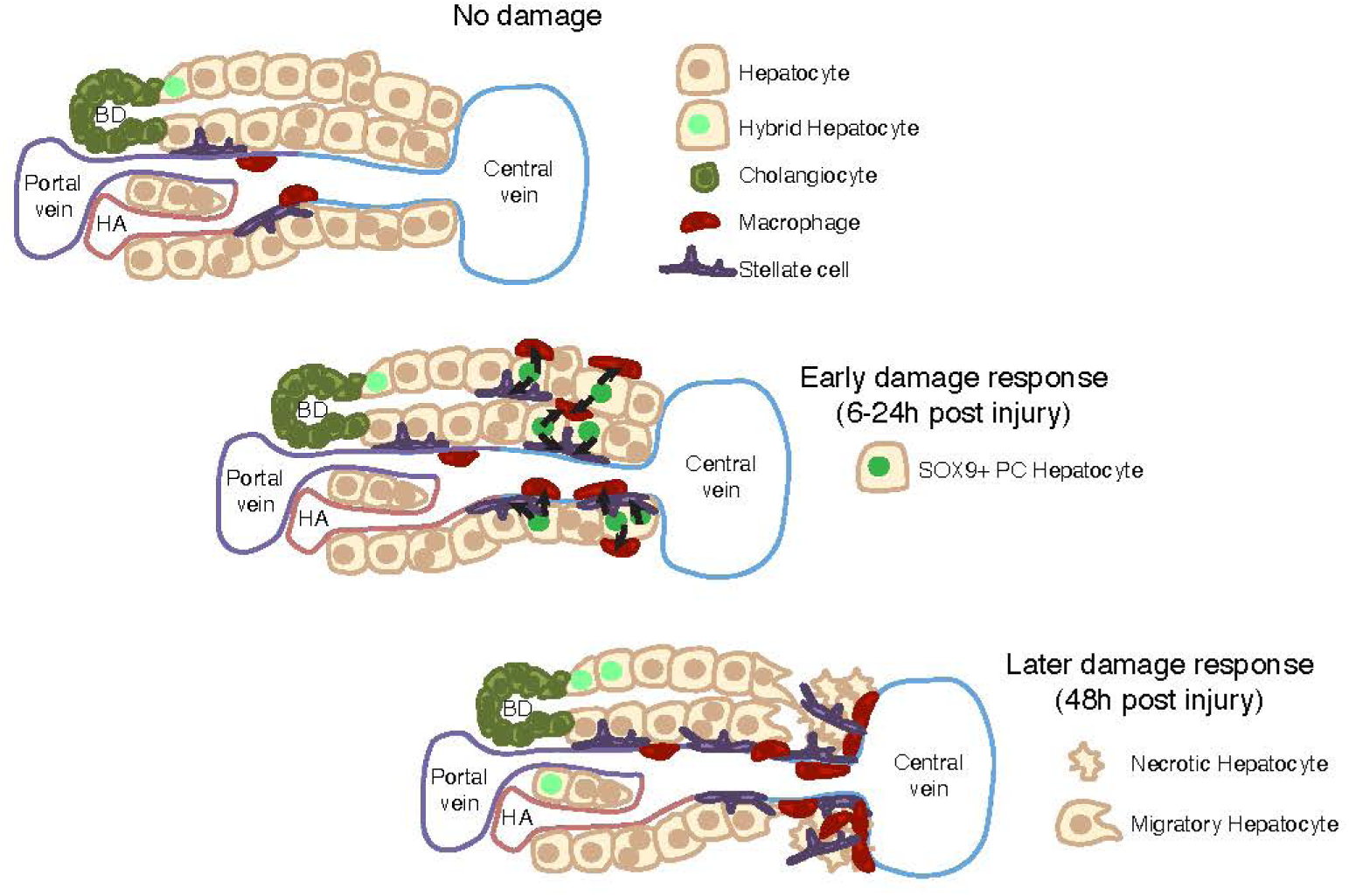
SOX9 mediates hepatocyte damage response at the early phase of acute liver injury. Schematic showing the liver lobule and the relevant cell types. Dynamics of SOX9 expression, and the infiltration of macrophages and HSCs are shown. Schematic adapted from (Gordillo et al., 2015).

Previous studies have demonstrated early SOX9 upregulation in hepatocytes in response to various forms of liver injury, including partial hepatectomy, ischemia reperfusion, and hepatic necrosis resulting from impaired intrahepatic bile duct formation during development (Antoniou et al., 2009; Qin et al., 2023; Shao et al., 2022; Swiderska-Syn et al., 2014). With the exception of ischemia-reperfusion injury, SOX9 expression in these models has been largely confined to periportal hepatocytes, and the functional significance of these SOX9-expressing cells remains poorly understood. Interestingly, Matchett and colleagues identified an ANXA2+ migratory hepatocyte population that is transcriptionally similar to our damage-responsive SOX9+ PC hepatocyte population (Figure S5a,b). Analysis of their publicly available snRNA-seq dataset revealed that *Anxa2*-expressing hepatocytes also show enrichment for *Sox9*, suggesting potential co-expression. Their ANXA2+ population emerged in response to human ALF, human chronic liver disease, and a range of mouse models of liver injury (pericentral, periportal, acute, and chronic) (Matchett et al., 2024). It is plausible that in these disease and injury contexts, ANXA2⁺ hepatocytes also co-express SOX9, although this remains to be formally demonstrated. Our findings thus support the emerging view that SOX9+ hepatocytes participate broadly in hepatic stress responses, integrating multiple aspects of the repair process across diverse injury contexts.

Intriguingly, studies have shown that hepatocyte-specific knock-out and knock-down of SOX9 during acute injury results in downregulation of cytokines such as TNFα, IL-6 and IL-1β (Fan et al., 2018; Qin et al., 2023). Data from Qin and colleagues further suggest that loss of hepatocyte-SOX9 impairs macrophage recruitment to sites of injury following CCl4-induced acute damage. These observations are consistent with our findings, which indicate that damage-responsive SOX9+ PC hepatocytes serve as important regulators of myeloid cell engagement during the early phase of acute liver injury. Moreover, several studies have demonstrated that disrupting macrophage recruitment or function delays tissue repair, prolongs necrosis, and exacerbates liver dysfunction in the context of acute injury (Hassan et al., 2023; Holt et al., 2010, 2008; Krenkel and Tacke, 2017; Strey et al., 2003; You et al., 2013; Zigmond et al., 2014). Thus, our findings highlight a previously unappreciated role for SOX9+ hepatocytes as a link between parenchymal stress responses and the orchestration of local myeloid cell dynamics.

While SOX9 expression has traditionally been associated with progenitor function, there is evidence pointing to damage-responsive roles for SOX9 in other regenerative contexts. For example, in the early phase of acute kidney injury (AKI), SOX9 plays a protective role by promoting the expression of pro-survival genes, thereby enhancing epithelial cell survival during injury (Kim et al., 2020). Similarly, in the mammalian rib, a *Sox9*-expressing periosteal subpopulation orchestrates large-scale bone regeneration. While these cells represent only a minority of the regenerating callus, they exert an essential non-cell-autonomous instructive role, stimulating neighboring cells to differentiate and contribute to bone repair (Kuwahara et al., 2019). These observations parallel our findings in the liver, where SOX9 potentially facilitates multiple cellular interactions that drive tissue repair, independent from progenitor-like properties of other SOX9+ cell populations.

Importantly, we observed that SOX9 expression in the damage-responsive PC hepatocytes is transient. In contrast, during AKI, renal tubular cells that fail to resolve repair maintain sustained SOX9 expression, a state associated with increased risk of progression to chronic kidney disease (Kumar et al., 2015). In addition to transient SOX9 expression (Figure 2h), our lineage tracing experiment showed a marked reduction in the RFP+ lineage-labeled cells by day 6 post-injury (Figure 3n), and RNA velocity analysis indicated an alternative trajectory for the Sox9-clusters that did not culminate in stable differentiated cell states (Figure 3q). These findings suggest that a substantial fraction of the damage-responsive SOX9+ PC hepatocytes may undergo cell death once their repair functions are completed, a mechanism that could be critical for restoring tissue homeostasis and preventing maladaptive remodeling leading to chronic disease.

Our data has also provided mechanistic insight into how SOX9 may perform its different functions during the early phase of acute injury. Epiregulon analysis suggests that SOX9 may promote the migratory hepatocyte phenotype by directly regulating *Anxa2*. In addition, this analysis also identified several upstream regulators of *Sox9*, including the the NF-κB family member *Rela* (NF-κB p65) (Table 8) (Ushita et al., 2009). Interestingly, many of these putative upstream regulators of *Sox9* were chromatin regulators and architectural proteins such as *Nipbl, Rad21, Smc1a, Brd4, Ncor1, Atrx,* and *Chd4* (Table 8), highlighting a potential role for dynamic chromatin remodeling in modulating SOX9 activity during the injury response.

In summary, our findings identify a transient, damage-responsive population of SOX9+ pericentral hepatocytes that is suggested to act as an early orchestrator of hepatic repair following acute injury. Intriguingly, by modulating myeloid cell recruitment and activity, SOX9+ hepatocytes appear to influence the broader inflammatory milieu critical for effective tissue repair. However, the exact effect on myeloid cell recruitment and activity upon hepatocyte-specific reduction/removal of *Sox9* was not uncovered in this study and remains to be determined. While these results advance our understanding of SOX9+ hepatocyte function in acute injury, further work will be needed to determine the relevance of these mechanisms across different injury models, and to assess whether they might be leveraged therapeutically to support regenerative outcomes.

## Methods

### Ethics Statement

Our research complies with all of the relevant ethical regulations. Animals were maintained in accordance with the Guide for the Care and Use of Laboratory Animals (National Research Council, 2011). Genentech is an Association for Assessment and Accreditation of Laboratory Animal Care-accredited facility, and all animal activities in this research study were conducted under protocols approved by the Genentech Institutional Animal Care and Use Committee.

### Animal strains and treatments

All animal experimentation was performed according to protocols approved by the Genentech Institutional Animal Care and Use Program (IACUC) committee. Female mice were used for all experiments. C57BL/6 mice were obtained from Charles River-Hollister, the transgenic *Sox9-CreER^T2^* mice were previously described (Kopp et al., 2011), and the inducible knock-in *Sox9-CreER^T2^* mice were generated in-house. For acute injury induction, 8-week-old mice were injected intraperitonially with a single sublethal dose of 0.5ml/kg CCl4 diluted in corn oil. For lineage tracing studies using the transgenic *Sox9-CreER^T2^* mice, a single dose of 200mg/kg of tamoxifen was administered through oral gavage at the indicated time points. For lineage tracing studies using the knock-in *Sox9-CreER^T2^* mice, a single dose of 5mg/kg of tamoxifen was administered intraperitonially at the indicated time points.

### Generation of mice carrying the *Sox9-Cre^ERT2^* knock-in allele

Mice carrying the *Sox9-Cre^ERT2^* allele were generated using established CRISPR-assisted ES cell methodology at Taconic Biosciences and C57BL/6NTac-derived ES cells. As the end result, a T2A-CRE^ERT2^ cassette was inserted right before the *Sox9* stop codon in exon 3, corresponding to the following genomic location: GRCm39/mm39 chr11:112,676,333. The sgRNA used had the following sequence: 5’-GCUUUUCUCUUCUCAGGGUC-3’ (reverse strand). Correctly targeted ES cell clones were identified using Southern blot and junction PCR analysis, followed by NGS-based analysis of 14 predicted potential CRISPR off-targets. Off-target-negative clones were injected to obtain chimeras. After germline transmission, the mouse allele was maintained on a C57BL/6N background.

The allele has been registered with MGI as Sox9^em1(cre/ERT2)Gne^.

### Histology and immunostaining

Mouse livers were collected at 6h, 24h, 48h and 6 days post damage induction. The left and median lobes were fixed with 10% neutral buffered formalin, processed and embedded in paraffin blocks, and then used to create 4 µm thick sections.

Hematoxylin and eosin (H&E) staining and IHC were performed by the Histopathology core lab at Genentech. IHC was performed using the following antibodies: anti-Ki-67 at 0.334 ug/ml (Cell Signaling Technologies, clone D3B5, 12202, rabbit), anti-SOX9 at 0.333 μg/mL (Millipore, AB5535, rabbit), and anti-RFP at 1 μg/ml (Rockland Immunochemicals for Research, 600-401-379, rabbit). All H&E and IHC slides were imaged on a NanoZoomer S360 brightfield whole-slide scanner (Hamamatsu, USA) at 20X.

Dual immunofluorescence for SOX9 and RFP was performed on the Ventana Discovery Ultra platform with CC1 Standard antigen retrieval at 97°C for 64 min. Sections were first incubated with anti-RFP at 1.1 μg/mL (Rockland Immunochemicals for Research, 600-401-379, rabbit). Following antibody denaturation with CC2 antigen retrieval at 100°C for 8 min, sections were incubated with anti-SOX9 at 0.5 μg/mL (Millipore, AB5535, rabbit). Slides were imaged on an Olympus VS200 at 20X.

Immunofluorescence was performed using the following antibodies: anti-HNF4a at 1:300 (R&D Systems, PP-H1415-00, mouse), anti-CYP2E1 at 1:100 (Abcam, ab28146, rabbit), anti-Ki67 at 1:200 (Thermo Fisher Scientific, SolA15 14-5698-82, rat), anti-SOX9 at 1:1000 (Millipore, AB5535, rabbit), anti-RFP at 1:1000 (Rockland Immunochemicals for Research, 600-401-379, rabbit), anti-Desmin at 1:100 (R&D Systems, AF3844, goat), anti-α-SMA-Cy3 (Milipore Sigma, C6198, mouse), and anti-F4/80 at 1:200 (Biorad, MCA497R, rat). Slides were imaged using the Leica Thunder microscope.

Dual immunofluorescence of same-host antibodies (SOX9 and CYP2E1, and RFP and CYP2E1) was performed through sequential staining. The first target was covalently labeled with Cy3, using Poly-HRP anti-rabbit (Leica Biosystems, PV6119) followed by TSA amplification (Akoya Biosciences, NEL744001). This was followed by a second heat-mediated antigen retrieval step to remove the antibodies. Standard immunofluorescence staining was subsequently used to detect the second target.

Slides were imaged using the Leica Thunder microscope. Images were analyzed using Fiji, and processed with Adobe Illustrator.

### Serum Biochemical assays

Blood samples were collected at 6h, 24h, 48h and 6 days post damage induction. The samples were centrifuged for 10 min at 5000 rpm, and kept at 4°C until analysis. Serum ALT levels were measured on the Beckman Coulter Au480 Clinical Chemistry Analyzer.

### RNAscope In Situ Hybridization

RNAscope in situ hybridization was performed on formalin-fixed, paraffin-embedded (FFPE) mouse liver sections using the RNAscope 2.5 HD Brown Assay Kit (Advanced Cell Diagnostics (ACD), Cat.No.322500), according to the manufacturer’s instructions. Briefly, 5 µm FFPE liver sections were baked at 60°C for 1 hour, followed by deparaffinization, hydrogen peroxide treatment, target retrieval, and protease digestion using reagents provided in the H_2_0_2_ & Protease Plus Reagents kit (ACD, Cat. No 322440). RNAscope probes targeting Sox9 (Mm-Sox9, ACD, Cat. No. 401051) or Cre recombinase (Cre, ACD, Cat. No. 312281) were hybridized to tissue sections for 2 hours at 40°C in a HybEZ oven. Signal amplification steps were performed as instructed in the manufacturer’s protocol, and hybridized probes were detected using the DAB chromogenic substrate provided it the kit, resulting in a brown precipitate at the site of probe binding. Slides were counterstained with 20% Gills’ Hematoxylin III (Fisher Scientific, Cat. No 353716), subjected to bluing with Scott’s water (Fisher Scientific, Cat. No.225000050), dehydrated, and mounted in Permount (Fisher Scientific, Cat. No SP15-100).

### Single-nucleus RNA and ATAC sequencing and computational analysis

#### Nuclei Isolation

Liver tissue (caudate lobe) from one oil-treated mouse and one CCl4-treated mouse was used for nuclei isolation. The tissue was frozen in liquid nitrogen upon harvest and stored at -80°C. Nuclei were isolated using the Chromium Nuclei Isolation Kit with RNase Inhibitor (10X Genomics, PN-1000494), and following the manufacturer’s protocol.

#### Library Prep and Sequencing

Nuclei were checked for quality using Cellometer K2 Image Cytometer cell counter Revvity (Waltham, MA), then tagmented, and then injected into microfluidic chips to form Gel Beads-in-Emulsion (GEMs) in the 10X Chromium X instrument (Pleasanton, CA). Following 10X Genomics protocol Chromium Single Cell Multiome ATAC + Gene Expression Reagent Kits User Guide (CG000338), Gene expression and ATAC libraries were generated and profiled using the Tapestation 4200 Agilent Technologies (Santa Clara, CA), and quantified with Qubit Fluorometer ThermoFisher Scientific (Waltham, MA). Gene Expression libraries were sequenced on a NovaSeq6000 Illumina (San Diego, CA) with a read format of 28 cycles for Read 1, 10 cycles for i7 index, 10 cycles for i5 index and 90 cycles for Read 2 with a targeted sequencing depth of 20,000 read pairs per cell. ATAC libraries were sequenced with a NextSeq 2000 lllumina (San Diego, CA) with a read format of 50 cycles for Read 1, 8 cycles for i7 index, 24 cycles for i5 index and 50 cycles for Read 2, with a targeted sequencing depth of 25,000 read pairs per cell. Raw sequencing data was demultiplexed by Illumina’s Bcl2Fastq software.

#### Data Processing

The demultiplexed data were processed using CellRanger ARC version 2.0.2, a software suite designed for single-cell multiome analysis. The reference genome used was the Mouse (GRCm38-2020-A), ensuring alignment and feature annotation were consistent with high-quality mouse genomic standards. During the processing, the gex_exclude_introns_input parameter was set to “False,” allowing the inclusion of intronic reads in the gene expression analysis, which can enhance the detection sensitivity of gene expression profiles.

#### Data Analysis

ArchR (Granja et al., 2021) version v1.0.1 was used to analyze chromatin accessibility and transcriptomic data.

#### snATAC-seq preprocessing

Arrow files were generated from fragment files using the ‘createArrowFiles’ function with the following parameters: minTSS = 4 and minFrags = 1000, with TileMatrix and GeneScoreMatrix enabled. We computed doublet scores using addDoubletScores (k = 10, LSIMethod = 1) and built an ArchRProject using the ‘ArchRProject’ function.

#### snRNA-seq preprocessing and integration

snRNA-seq count matrices were imported using ‘import10xFeatureMatrix’. A shared set of barcodes was identified between the snRNA-seq and snATAC-seq datasets, and both modalities were subset accordingly. The shared snRNA-seq data were added to the ArchR project using ‘addGeneExpressionMatrix’.

#### Quality control

Doublets were filtered out using ‘filterDoublets’ with the parameters ‘cutEnrich = 1’, ‘cutScore = -Inf’, and ‘filterRatio = 1’. Cells without any RNA UMIs, and a mitochondrial gene expression ratio of ≥ 0.25 were also filtered out.

#### Dimensionality reduction and clustering

Dimensional reduction was performed at the ‘TileMatrix’ and ‘GeneExpressionMatrix’ levels using ‘addIterativeLSI’. For gene expression data dimension reduction, the ‘depthcol’ parameter was set to ‘Gex_nUMI’. Harmony batch-corrected reduced dimensionality results were then obtained using the ‘addHarmony’ function. Subsequently, the batch-corrected dimensionality reduction results from the two modalities were combined using ‘addCombineDims’. Clustering was performed independently on both the individual modality datasets and the combined results with ‘addClusters’. UMAPs were added in a similar manner using ‘addUMAP’.

The same approach was applied on the hepatocyte subset, and the damaged sample subset (no Harmony added for the damaged sample only). For the hepatocyte subset, hepatocyte-specific clusters were extracted, retaining nuclei only from these clusters. For the damaged subset, the ArchR project was subset by “Sample”.

#### Construction of SingleCellExperiment Object

Gene expression counts were extracted from the ArchR *GeneExpressionMatrix* and converted to a SingleCellExperiment (SCE) object using ‘SingleCellExperiment’ (Amezquita et al., 2020). Gene annotations and UMAP coordinates were appended to the SCE object. Counts were log-normalized using ‘logNormCounts’ from the scuttle package (McCarthy et al., 2017).

#### Marker detection and pathway enrichment

Marker genes for all clusters were identified using ‘getMarkerFeatures’ (Wilcoxon test, bias-adjusted for nUMI). Pathway enrichment was performed using ‘GSEA’ from the clusterProfiler package (Yu et al., 2012), with MSigDB gene sets.

#### Adding Module Scores

The GeneExpressionMatrix SCE was converted into a Seurat object (using ‘as.Seurat’) (Seurat v5.1.0) (Hao et al., 2021) for adding module scores, using ‘AddModuleScore’. The central and portal zonation scores were scaled to a value between 0 and 1, and then the zonation specificity scores were calculated using the following formula: zonation score = central score/(central score + portal score) (Matchett et al., 2024).

#### Visualization

UMAP and dot plot visualizations were generated using the dittoSeq package (Bunis et al., 2020).

#### Peak calling and Motif enrichment

Pseudo-bulk profiles were generated using ‘addGroupCoverages’, and chromatin accessibility peaks were identified using MACS2 via ‘addReproduciblePeakSet’. A PeakMatrix was created, and marker peaks for all clusters were identified using ‘getMarkerFeatures’. Peaks were annotated with gene information using ‘addPeakAnnotations’, and motif enrichment was performed with ‘peakAnnoEnrichment’.

#### Transcription factor activity

Transcription factor (TF) activity was inferred using the Epiregulon v1.1.6 package (Włodarczyk et al., 2024). The process involved several steps: First, accessible chromatin peaks (regulatory elements – REs) were linked to their putative target genes (TGs) using ‘calculateP2G’, if their chromatin accessibility and gene expressions were highly correlated across cell clusters (correlation coefficient > 0.5). Next, TF motif binding information was added by overlapping the regions of the peak matrix with the ChIP-seq database, using ‘addTFMotifInfo’. Initial TF-RE-TG triplet networks were constructed using ‘getRegulon’, and pruned using the chi-square test of independence (‘pruneRegulon’), retaining only triplets where TF expression, RE accessibility, and TG expression co-occur more frequently than expected by chance. The strength of regulation was estimated using the Wilcoxon test (‘addWeights’). This test compared target gene expression in cells jointly expressing all three elements (TF-RE-TG) versus cells that did not. The activity of a transcription factor was computed as the weighted sum of all its target genes, using ‘calculateActivity’. Finally, TFs with cluster-specific activity were identified using ‘findDifferentialActivity’.

#### Cell-Cell Communication Analysis using CellChat

Ligand-receptor communication networks among cell populations in the damaged liver sample were inferred using CellChat v1.0.0 (Jin et al., 2021). A CellChat object was created using ‘createCellChat’, with the SCE log-normalized counts used as input and the cluster identities used to group cells. The built-in CellChatDB.mouse ligand-receptor interaction database was employed to define candidate interactions. The data were preprocessed to identify over-expressed genes and over-expressed interactions using the functions ‘identifyOverExpressedGenes’ and ‘identifyOverExpressedInteractions’. Communication probabilities between cell populations were then computed using ‘computeCommunProb’, with the truncated mean method (trim = 0.1). Only interactions supported by at least 10 cells per group were retained via ‘filterCommunication’. We further inferred signaling pathway-level communication using ‘computeCommunProbPathway’, and summarized aggregated communication networks with ‘aggregateNet’. Visualization of the resulting communication networks was performed using netVisual_circle(), enabling visualization of interaction strengths (weight) among clusters.

#### Data conversion for Python-based analysis pipelines

The GeneExpressionMatrix SCE was converted into a H5AD file using the zellkonverter v1.15.4 R package (DOI: <u>10.18129/B9.bioc.zellkonverter</u>) for analysis in Python. The H5AD file was created into an AnnData object using scanpy v1.11.2 (Wolf et al., 2018).

#### Cell-Cell Communication Analysis using LIANA

LIANA v1.5.1 (Dimitrov et al., 2022) was also used for inferring ligand-receptor interactions among cell populations in the damaged liver sample. By default, LIANA uses human gene symbols. These were first homology mapped to mouse symbols. To obtain a robust consensus ranking of predicted cell-cell communication events, the Rank Aggregate approach was employed. Aggregate ranks were used for downstream visualization and interpretation of the cell-cell communication landscape in our dataset.

#### RNA velocity Analysis

RNA velocity analysis was performed to infer the future transcriptional states of single cells based on spliced and unspliced mRNA abundance using Velocyto v0.17.17 and Scvelo v0.3.2 (Manno et al., 2018). Velocyto was used to generate loom files, using the GRCm38-2020-A/mm10 annotation and repeat mask. The AnnData and Loom objects were merged using ‘scv.utils.merge’ to obtain a combined object with both RNA velocity and gene expression information. Quality control, filtering, and normalization were performed using ‘scv.pp.filter_and_normalize’. Principal component analysis (PCA) and neighborhood graph construction were performed with ‘sc.pp.pca’ and ‘sc.pp.neighbors’. Moments for velocity estimation were computed with scv.pp.moments’. RNA velocity dynamics were modeled with ‘scv.tl.recover_dynamics’, followed by estimation of velocity vectors using ‘scv.tl.velocity’ and ‘scv.tl.velocity_graph’. The resulting velocity streams projected onto UMAP embeddings were visualized using ‘scv.pl.velocity_embedding_stream’. To identify root and terminal cell states, ‘scv.tl.terminal_states’ was used.

### GeoMx and computational analysis

#### Sectioning and ROI pre-selection

Formalin-fixed paraffin-embedded (FFPE) blocks were sectioned at 4 µm and processed using the GeoMx protocol for RNA assays. Sections serial to the GeoMx collection slides were stained with anti-CYP2E1 primary antibody (PA5-79131, Invitrogen) at 0.25 µg/mL for 60 mins at room temperature. An Alexa Fluor 647-conjugated Donkey anti-Rabbit secondary antibody was applied next at 4 µg/mL for 30 mins at room temperature followed by DAPI counterstain (Invitrogen, cat# D1306) for 16 minutes at 0.2 µg/ml. The slides were mounted with ProLong Gold Antifade Mountant (Invitrogen, cat# P36930) and imaged with the Olympus VS200 Research Slide Scanner. The study pathologist selected ROIs from these images with polygonal annotations, keeping within the maximum collection area of 660 x 780 µm.

#### GeoMx Slide Prep

GeoMx collection slides were prepared for collection according to Nanostring’s guidelines for RNA NGS assays. Briefly, slides were baked at 70°C for 30 minutes and moved to the Leica Bond RX for deparaffinization and post fixation. Antigen retrieval was performed using ER2 (AR9640, Leica) at 100°C for 20 minutes, followed by proteinase K digestion (AM2546, ThermoFisher) at 1µg/mL for 15 minutes. Post fixation was performed using 10% neutral buffered formalin (16004-122, VWR). Slides were removed from the Bond RX and incubated overnight (18 hours) at 37°C in a hybridization oven (HybEZ, ACD) with RNA probes from GeoMx’s Human Whole Transcriptome Atlas. The next day, slides were washed twice in a stringent solution at 37°C for 25 minutes per wash. Slides were blocked with Nanostring’s Buffer W (blocking buffer) for 30 mins. Anti-CYP2E1 primary antibody (PA5-79131, Invitrogen) was applied at 0.25 µg/mL for 60 mins at room temperature. Syto13 counterstain (57575, Invitrogen) was included in the primary antibody mix at 500 nM. An Alexa Fluor 647-conjugated Donkey anti-Rabbit secondary antibody was applied next at 4 µg/mL for 30 mins at room temperature. Following visualization marker incubation, slides were washed twice with 2× saline sodium citrate (SSC) for 5 minutes and then loaded onto the GeoMx instrument.

#### Library Prep and QC

The annotated serial section images were used to guide region of interest (ROI) selection. Polygonal ROIs were selected and photocleaved oligonucleotides were collected in a 96-well plate. Sample index PCR and SPRIselect-mediated cleanup (Beckman Coulter #B23319) were performed to produce the next-generation sequencing (NGS) library following Nanostring’s guidelines and using GeoMx’s Seq Code Pack. Library concentration was determined using a Qubit 4 Fluorometer (Invitrogen, Waltham, MA). Library quality was assessed using an Agilent TapeStation 4200 with D5000 dsDNA reagents (Agilent #5067-5588/5589). FASTQ files were processed through the “geomxngspipeline” program, version 2.3.3.10 (Nanostring), using default parameters.

#### GeoMx computational analysis

GeoMx data were processed using the *GeomxTools* R package. In short, raw DCC files were read and subjected to segment level and probe level QC, followed by collapsing probes onto the gene level, segment gene detection based on limit of quantification, gene detection rate filtering for genes across the study and finally quartile (Q3) normalization as described in the package *GeomxTools* R package vignette. A mixed model analysis, considering the slide as a random effect, was performed using the *mixedModelDE* function in the *GeomxTools* R package to identify differentially expressed genes between damaged and undamaged regions of interest. Covariance analysis within damaged samples was performed using the *WGCNA* package in R for the top 1000 most variable genes, a soft thresholding power of 12 for network construction and dynamic treecutting using a Topologic Overlap Matrix calculated based on bidweight midcorrelation to determine adjacencies. We tested for the enrichment of particular gene categories to identify relevant biological processes associated with (1) differential expression between damaged and undamaged ROIs or (2) covariance modules, by using functions from the “clusterProfiler” R package (Wu et al., 2021) and the MSigDB gene set collection version 7.2 (Liberzon et al., 2015). In the case of differential expression, damaged ROIs were contrasted with undamaged ROIs, genes were ranked based on the π statistic (π=logFC * -log10(p-val)) (Xiao et al., 2014) and subsequently gene set enrichment was performed using the clusterProfiler::GSEA function. In the case of covariance modules, genes associated with specific modules were evaluated using a hypergeometric test using the clusterProfiler::enricher function.

## Supporting information

All Supplemental Figures

## Acknowledgements

We are grateful to C. Cottonham, S. Hankeova, G. Hernandez, and E. Reyes for helpful discussions. We thank the Genentech Research Pathology, Necropsy, Histology, and NGS laboratories for their experimental contributions, and the genetically engineered mouse models lab for construction of the knock-in *Sox9-CreER^T2^* mouse line.

## Author contributions

S.N.F.A. designed the project, performed experiments, data analysis, the sn-multiome computational analysis, and wrote the manuscript. K.S. performed all experiments related to the transgenic *Sox9-CreER^T2^* line. G.L. advised on initial analysis for the multiome data. M.F. performed all experiments for GeoMx. R.P. performed the computational analysis for the GeoMx data. E.R.S. provided histopathological analysis and discussion. C.W.S. and L.V. directed and supervised the research, and analyzed the data. L.V. also contributed to writing the manuscript. All authors approve the content of the manuscript.

## Declaration of interests

All authors are or where employees of Genentech Inc. and/or hold shares in Roche. C.W.S is currently employed at Gilead Sciences.

---

**Figure S1. Early stages of damage and repair progression in CCl4-induced acute injury. (Related to Figure 1)**

(a-e) Representative H&E images from mice treated with oil (a) or CCl4, 6h (b), 24h (c), 48h (d), and 6d (e) post treatment (n = 3 for oil, and n = 5 for CCl4, for each time point). CV = central vein, PV = portal vein. Arrows represent swollen hepatocytes. (f,g) Representative co-stain images for HNF4a and CYP2E1 in mice treated with oil (f), and CCl4 at 6d (g) post treatment. (h) Representative co-stain image for Ki67 and CYP2E1 in mice treated with oil. (i) Quantification of the percentage of proliferating hepatocytes in the central or the portal region after oil or CCl4 treatment, at the indicated time points. Comparison between oil and CCl4 treatment – p-values: 6h, central = 0.95, portal = 0.39, 24h, central = 0.91, portal = 0.89, 48h, central = 1.34E-09, portal = 4.34E-13. (j) Representative IHC images for Ki67 at 6d for oil and CCl4 treatment. Student’s T-test used for statistical analysis. Error bars represent SEM. Scale bars are 100 μm.

**Figure S2. Single nuclear transcriptomics analysis at 24h after injury. (Related to Figure 1)**

(a) Quality control metrics (number of genes, and mitochondrial percentage) from both undamaged (oil, n = 1) and damaged (CCl4, n = 1) sample. (b) UMAP showing the clusters obtained for all cells from both oil and CCl4 samples. (c) UMAP showing the clusters split by sample. (d) Dotplot annotating clusters by lineage signature expression (Table 1). (e) UMAP of cell lineages based on known lineage signatures (Table 1). (f) UMAP of portal and central hepatocyte scores (Table 1). (g) UMAP of hepatocyte subset showing new subclusters, split by sample. (h) Overlay of the cell lineages from (e) onto the hepatocyte subset, split by sample. (i) Hepatocyte signature (Table 1) for the hepatocyte subset, split by sample.

**Figure S3. Discovery of a novel SOX9+ cell population activated during early acute injury. (Related to Figure 2)**

(a) Representative co-stain image for HNF4a and SOX9 in an undamaged liver. Arrow represents a hybrid hepatocyte. (b) Representative co-stain image for SOX9 and CYP2E1 in mice treated with CCl4 at 48h post treatment. (c) Representative IHC images for SOX9 at 6d for oil and CCl4 treatment. (d) Quantification of the percentage of SOX9+ nuclei outside the CYP2E1 boundary (in periportal hepatocytes) at 48h. p-value = 1.26E-09. (e-g) Representative co-stain images for SOX9 and Ki67 at 48h. Arrows represent SOX9+/Ki67+ hepatocytes, arrowheads represent SOX9+ only hepatocytes. (h) Quantification of the percentage of proliferating SOX9+ hepatocytes at 48h. p-value = 3.31E-10. (i) UMAP showing *Sox9* expression for both oil and CCl4 samples. Student’s T-test used for statistical analysis. Error bars represent SEM. Scale bars are 100 μm.

**Figure S4. Lineage tracing the damage-responsive *Sox9+* cells reveals non-progenitor behavior. (Related to Figure 3)**

(a) Schematic of the transgenic *Sox9-CreER^T2^* mouse line. (b) Representative IF images for RFP, with tamoxifen (tam) given 18h prior to CCl4 injection. Tissue harvested at 24h and 6d post CCl4 injection. (c) Representative IF image for RFP, with tam administration 16h prior to CCl4 injection. Tissue harvested at 6h post CCl4 injection. (d) Representative IF image for RFP, with tam administration 3.5h prior to CCl4 injection. Tissue harvested at 16h post CCl4 injection. (e) Representative RNA Scope image for *Sox9* mRNA, with tam administration 16h prior to CCl4 injection. Tissue harvested at 6h post CCl4 injection. (f) Representative RNA Scope image for *Cre* mRNA, with tam administration 16h prior to CCl4 injection. Tissue harvested at 6h post CCl4 injection. (g) Schematic illustrating the generation of the knock-in *Sox9-CreER^T2^*mouse line. (h) Representative IHC image for RFP, with tam administration in the knock-in *Sox9-CreER^T2^* mice, without damage induction. Tissue harvested 7d post tam administration. (i,j) Representative co-stain images for RFP and SOX9, with tam administration 24h prior to oil (i) and CCl4 (j) injection. Tissue harvested at 24h post treatment. (k,l) Representative co-stain images for RFP and SOX9, with tam administration 5h post oil (k) and CCl4 (l) injection. Tissue harvested at 6d post treatment. (m) Quantification of the percentage of RFP+ hepatocytes in the central and the portal regions at the indicated time points. Comparison between oil and CCl4 for 24h – p-values: central = 3.01E-16, portal = 2.08E-22, for 48h – p-values: central = 2.20E-17, portal = 4.75E-19, for 6d – p-values: central = 6.31E-08, portal = 6.12E-11. (n) RNA velocity analysis of all the clusters in the damaged sample. Student’s T-test used for statistical analysis. Error bars represent SEM. Scale bars are 100 μm.

**Figure S5. *Sox9+* cells potentially mediate repair by interacting with stellate cells. (Related to Figure 4)**

(a) UMAP showing *Anxa2* expression for the hepatocyte subset, split by oil and CCl4 samples. (b) UMAP showing Migratory hepatocyte signature (Table 4) for the hepatocyte subset, split by oil and CCl4 samples. (c-g) Representative co-stain images for Desmin and α-SMA in mice treated with oil (c) or CCl4 at 6h (d), 24h (e), 48h (f), and 6d (g) post treatment. (h-j) Representative co-stain images for Desmin and SOX9 in mice treated with oil (h) or CCl4 (I,j), 24h post treatment. Scale bars are 100 μm.

**Figure S6. *Sox9+* cells display immune signatures and interact with macrophages at the site of injuy. (Related to Figure 5)**

(a) Representative IF images for CYP2E1, 24h after oil (undamaged, n = 3) or CCl4 (damaged, n = 5) treatment. Representative ROIs used for GeoMx sequencing are also shown. Undamaged ROIs = 30, damaged ROIs = 65. Scale bars are 1000 μm. (b) Representative ROIs from undamaged and damaged conditions shown at a higher magnification. Scale bars are 500 μm. (c) PCA of all undamaged and damaged ROIs. (d) *Cyp2e1* and *Sox9* gene expression for all undamaged and damaged ROIs. (e) Reactome Pathways Enrichment for the damaged ROIs (compared to the undamaged ROIs). (f-j) Representative IF images for F4/80 in mice treated with oil (f) or CCl4 at 6h (g), 24h (h), 48h (i), and 6d (j) post treatment. Scale bars are 100 μm.

