## Supplementary material for "Acute Pericentral Liver Injury Induces a Novel Transient Hepatocyte Population": All Supplemental Figures

**Supplemental Figure 1. Early stages of damage and repair progression in CCl4-induced acute injury.**

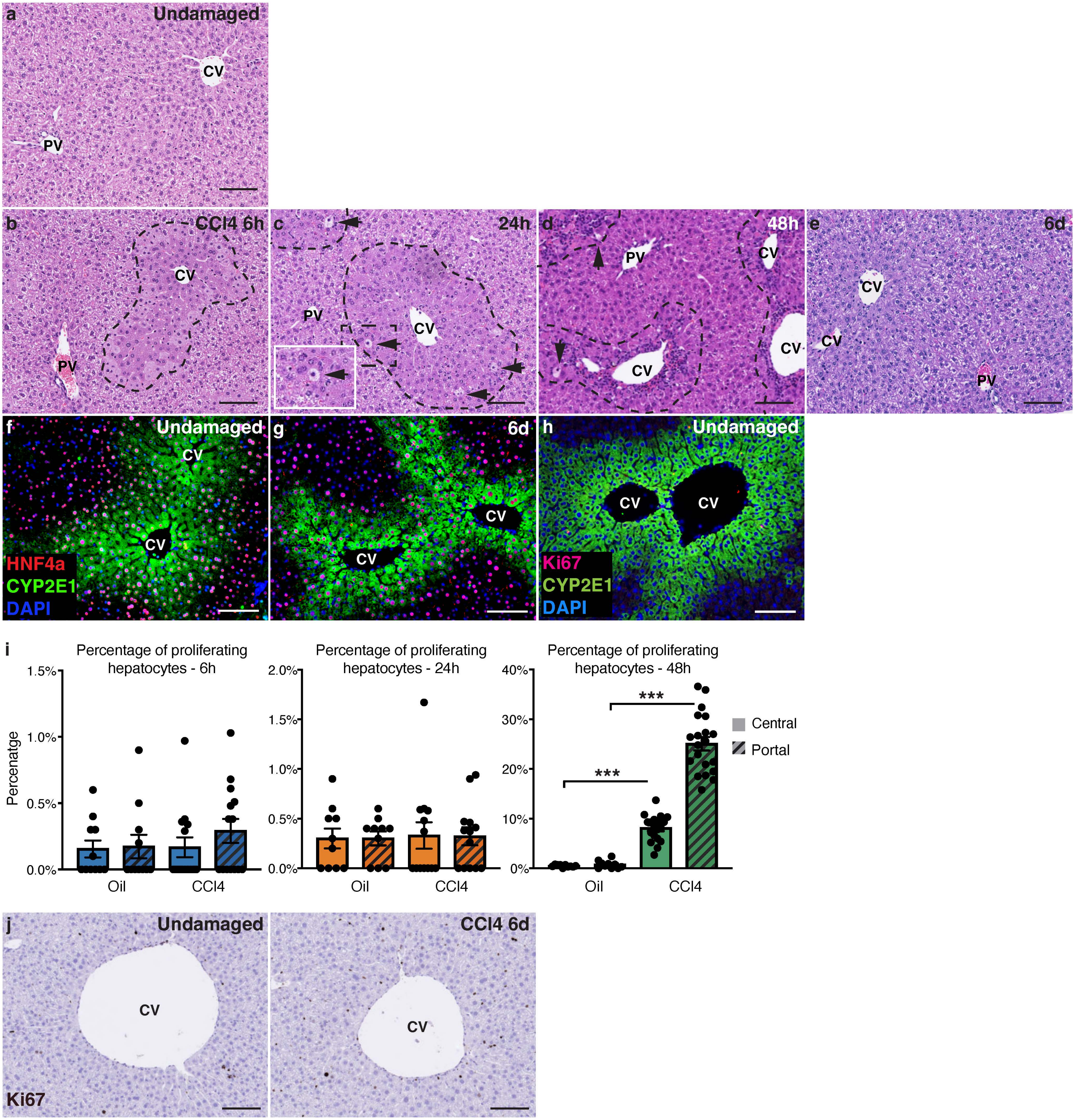

**Supplemental Figure 2. Single nuclear transcriptomics analysis at 24h after injury.**

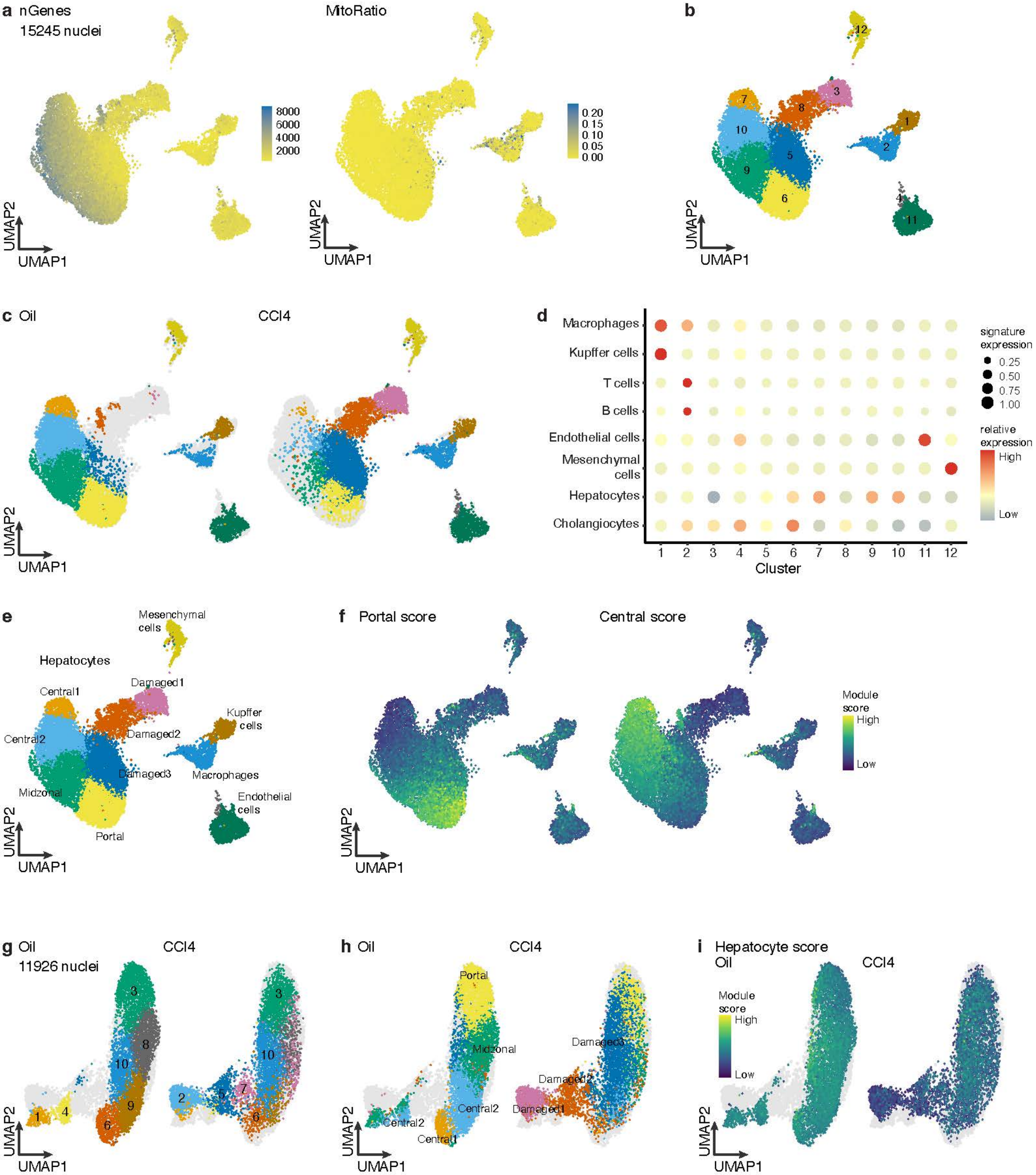

**Supplemental Figure 3. Discovery of a novel SOX9+ cell population activated during early acute injury.**

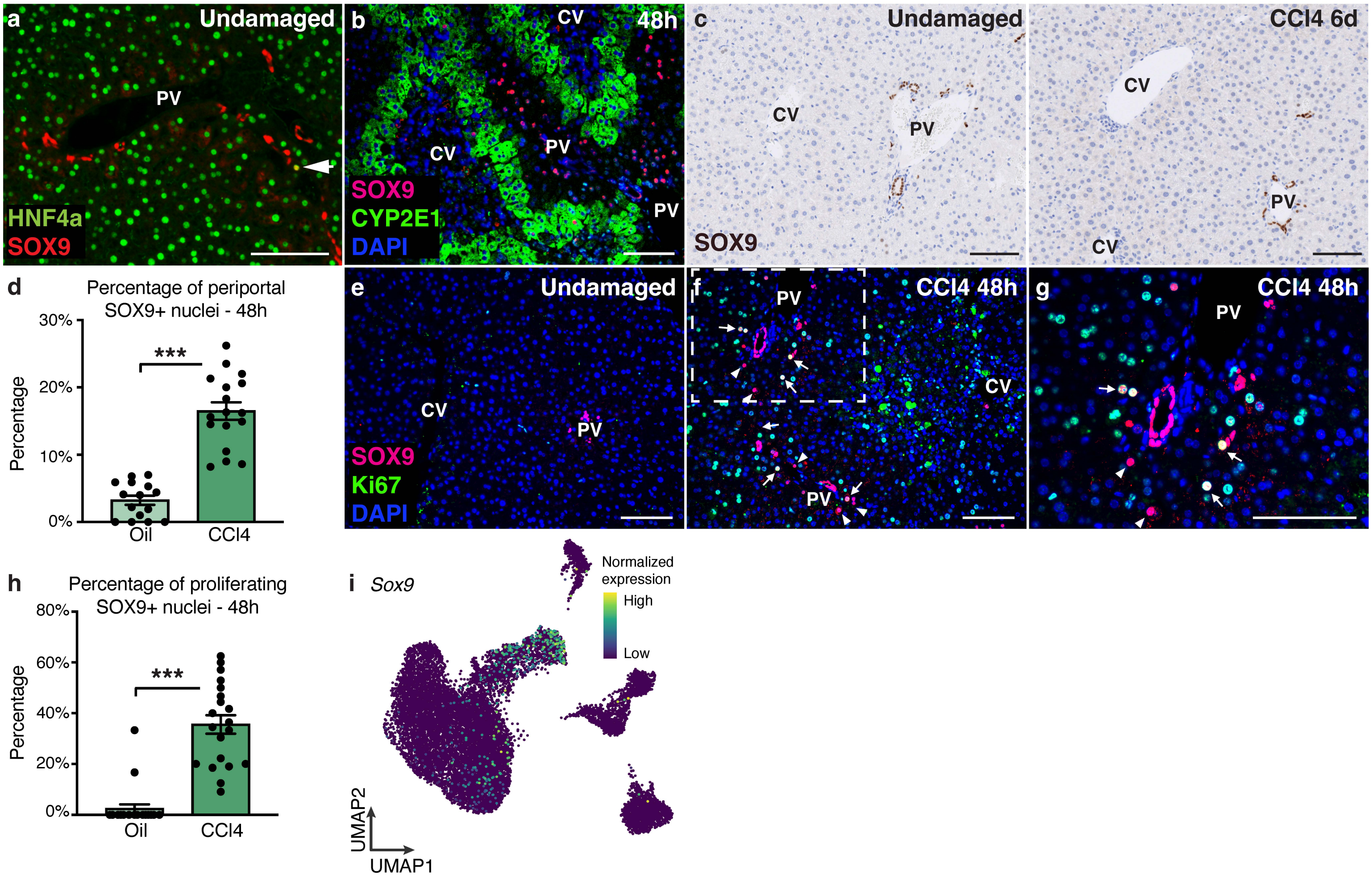

**Supplemental Figure 4. Lineage tracing the damage-responsive *Sox9+* cells reveals non-progenitor behavior.**

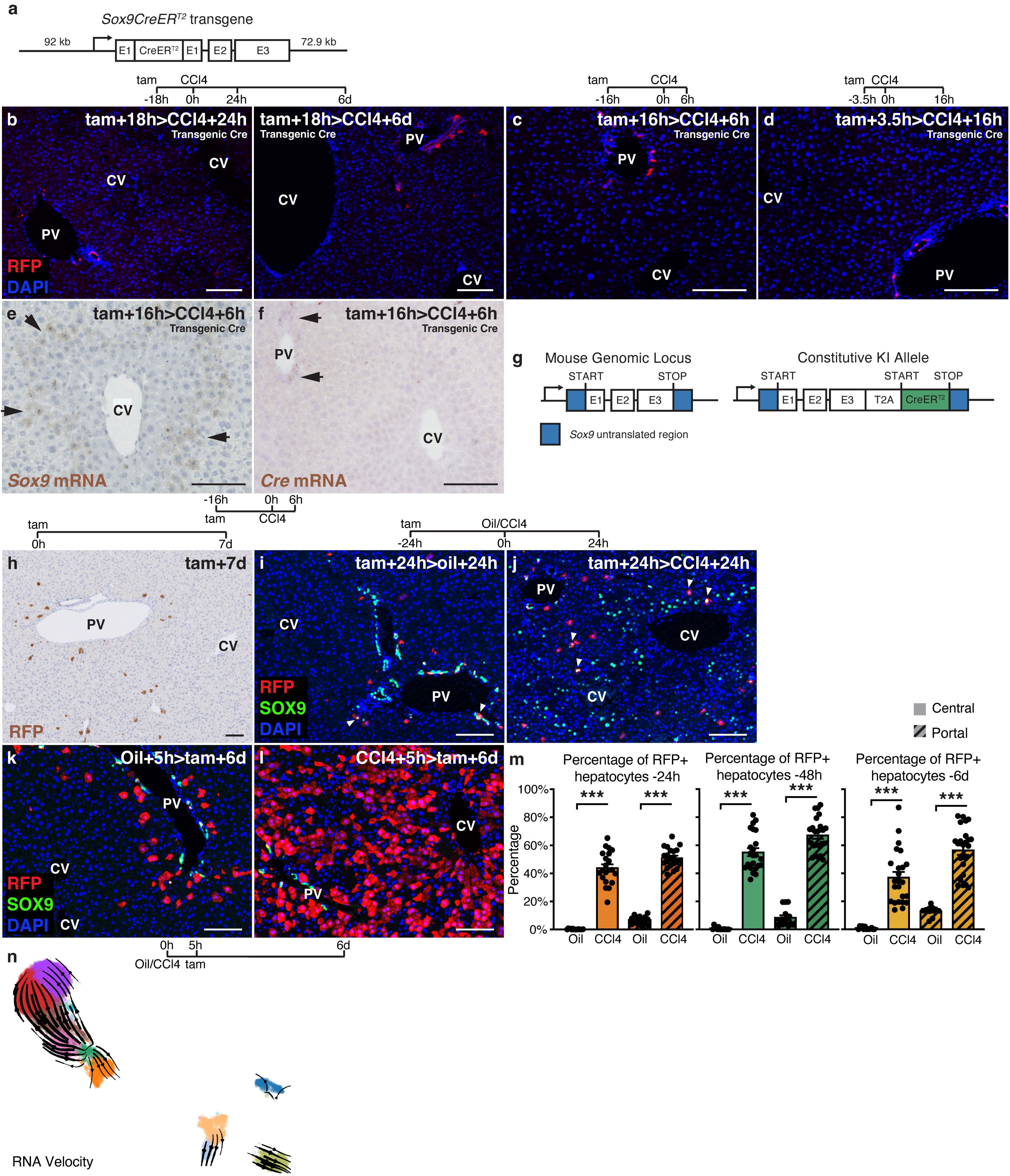

**Supplemental Figure 5. *Sox9+* cells potentially mediate repair by interacting with stellate cells.**

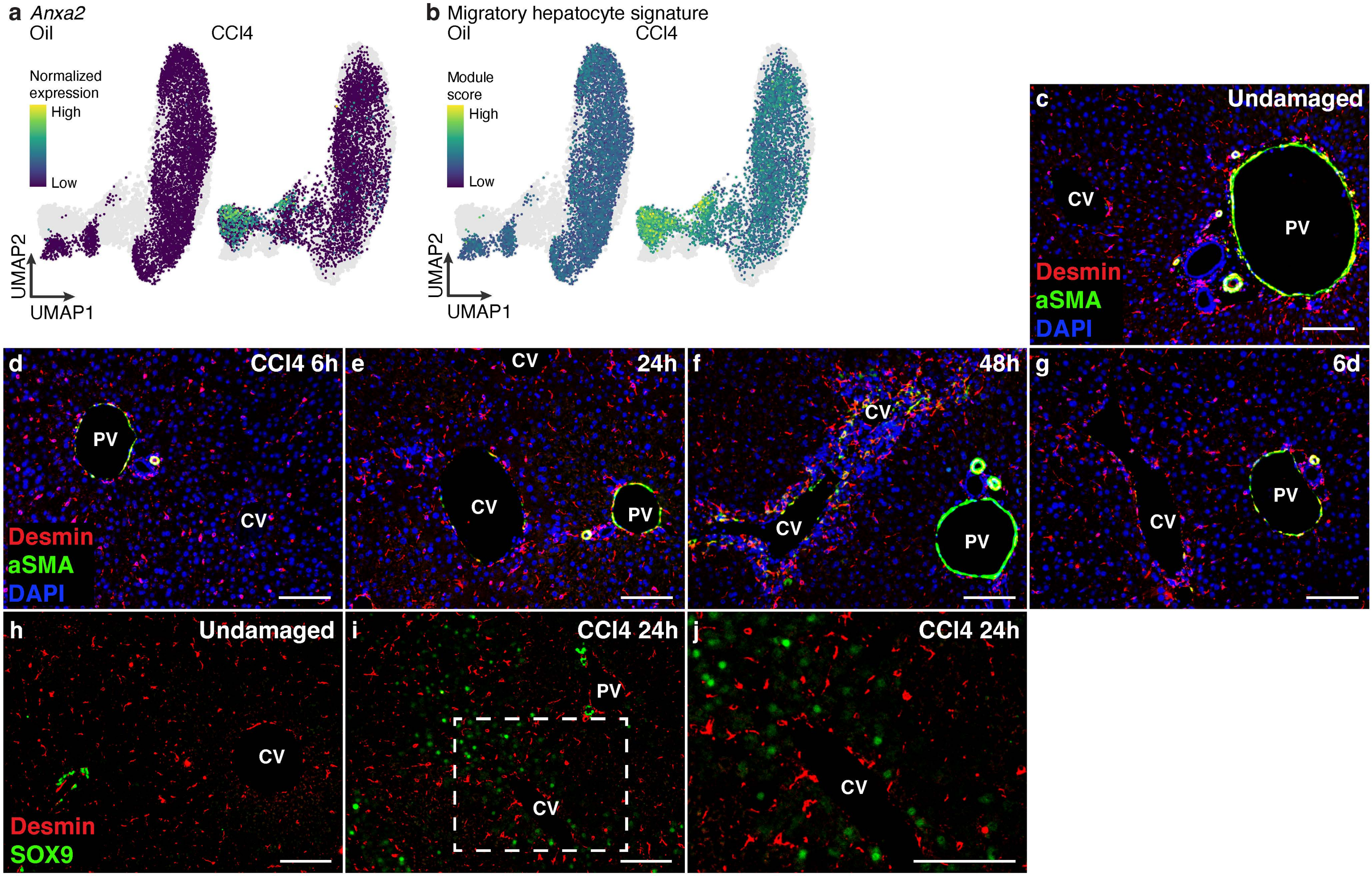

**Supplemental Figure 6. *Sox9+* cells display immune signatures and interact with macrophages at the site of injury.**

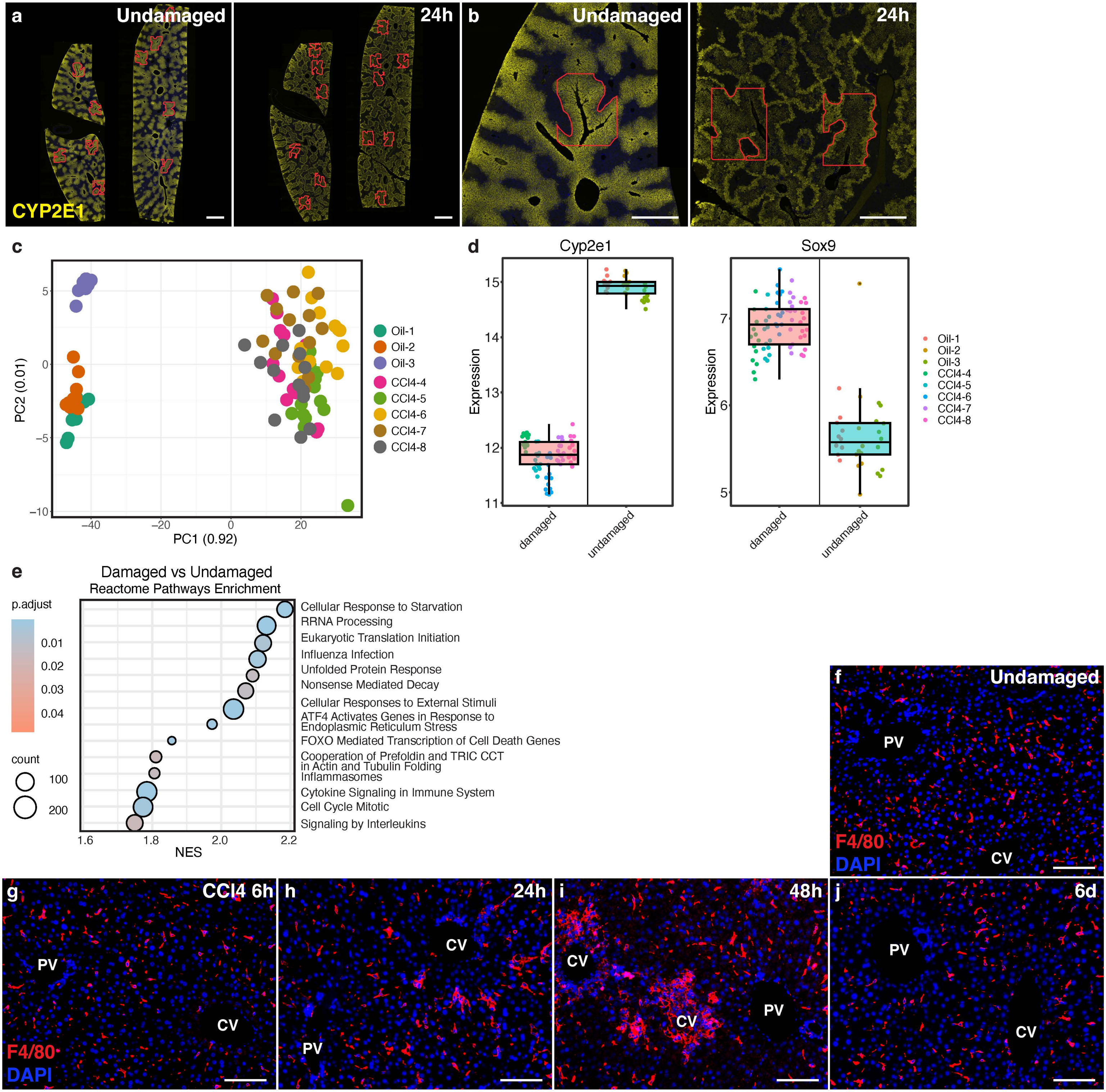
